# Participant followup rate can bias structural imaging measures in longitudinal studies

**DOI:** 10.1101/2021.02.10.430674

**Authors:** Richard Beare, Gareth Ball, Joseph Yuan-Mou Yang, Chris Moran, Velandai Srikanth, Marc Seal, the Alzheimer’s Disease Neuroimaging Initiative

## Abstract

Longitudinal MRI analysis is essential to accurately describe neuroanatomical changes over time. Loss of participants to followup (dropout) in longitudinal studies is inevitable and can lead to great difficulty in interpretation of statistical results if dropout is correlated with a study outcome or exposure. Beyond this, technical aspects of longitudinal MRI analysis require specialised processing pipelines to improve reliability while avoiding bias towards individual timepoints. In this article we test whether there is an additional problem that must be considered in longitudinal imaging studies, namely whether dropout has an impact on the function of FreeSurfer, a popular software pipeline used to estimate important structural brain metrics.

We find that the number of acquisitions available per individual can impact the estimation of cortical thickness and brain volume using the FreeSurfer longitudinal pipeline, and can induce group differences in brain metrics. The effect on trajectories of brain metrics is smaller than the effect on brain metrics.

**Highlights:**

- Longitudinal MRI analysis is essential to accurately track neuroanatomical changes over time
- Longitudinal MRI analysis requires specialised processing pipelines to reduce bias towards single timepoints
- Participant drop out or loss can bias neuroanatomical measures derived from longitudinal pipelines
- We find that group differences in the number of acquisitions available to analyse can cause group differences in estimated cortical thickness and brain volume
- This bias appears to be due to the number of scans used to create individualised templates in the Freesurfer longitudinal pipeline
- The effect on estimates of brain metric trajectories appears smaller than the effect on the estimates of brain metrics

## 1 Introduction

Brain metrics derived from magnetic resonance (MR) imaging in longitudinal studies are an increasingly important part of studies of development, aging and disease. In this article we investigate if the number of acquisitions available per participant to a longitudinal MR image processing pipeline has a measurable effect, at the group level, on estimates of cortical thickness, brain volume and change in these metrics over time. This effect, if present, differs from the well established survivor bias problems encountered in longitudinal studies in that it leads to a change in the *value* used in the statistical analysis. Survivor bias, on the other hand, may lead to incorrect assumptions about the study populations if not properly analysed. Such changes in estimated values potentially complicate the interpretation of longitudinal differences in some MR metrics even further.

Bias in followup rates can make analysis and interpretation of data collected by longitudinal studies extremely difficult, especially when participant dropout is correlated with a study outcome or exposure in an indirect or unexpected way. Survivor bias is a well known problem - for example smokers are more likely to be lost to followup in longitudinal studies of aging than non-smokers and thus great care is needed when interpreting differences in rates of disease, or other measures, in those remaining in the study. Demographic, socio-economic and other risk factors that are relevant to study questions, can contribute to risk of loss of follow up in many other ways. The traditionally recognised problems with bias in followup rate relate to *selection bias*, meaning that populations are not randomised.

The nature of bias examined in this article differs - it influences the raw numeric estimates produced by an MR image processing pipeline. Specifically, we examine whether there is a detectable difference between two sets of brain metrics, one estimated using *i* scans per person and the other using *j* scans per person, derived from one group of people. Group level differences that are attributable to differences between *i* and *j* have the potential to exaggerate or hide effects of interest in longitudinal studies.

The specific MR metric investigated in this analysis is the mean cortical thickness estimate generated by the FreeSurfer suite (Dale, Fischl, and Sereno 1999; Fischl and Dale 2000). FreeSurfer is a widely used neuroimaging package (freely available from http://surfer.nmr.mgh.harvard.edu/) capable of many forms of analysis and one of the core functions it performs is segmentation and characterization, including vertex-wise thickness estimation, of the cortex. FreeSurfer also includes a longitudinal processing pipeline (Reuter et al. 2012) in which all acquisitions for a study participant are combined to create a participant-specific template that contributes to the estimation of cortical thickness from each acquisition. Great care has been taken in the development of the longitudinal pipeline to ensure that the procedure is not biased by any of the acquisitions. The participant-specific template approach improves processing robustness in the presence of artifacts that might be present in some acquisitions, but is a potential source of difference at the study level via the number of acquisitions contributing to the template, which will be referred to as “*template depth*” in this article. There is potential for such biases to lead to over or under estimation of group effects if the number of acquisitions is correlated with group membership and the number of acquisitions influences the thickness estimate.

This study explores the issue of template depth causing biases in cortical thickness and supratentorial volume estimates produced by the longitudinal FreeSurfer pipeline, and trajectories of both, via simulated atrophy of scans from the Alzheimer’s Disease Neuroimaging Initiative (ADNI).

## 2 Code and data availability

All data used in this article is derived from the ADNI study and may be obtained from the referenced sources. Tools used to generate and process images and perform statistical analysis are publicly available and details are provided in Methods, below.

## 3 Methods

### 3.1 Study Design

A series of repeated measures experiments using MR scans with constant rates of simulated atrophy were used to explore the study questions. Each experiment assessed brain metrics derived from two or more groups consisting of the same individuals. The number of acquisitions per individual differed by group. Differences between groups would therefore be expected to be Type 1 errors or result from bias in estimation of brain metrics. Experiments involving estimates produced by the cross-sectional processing pipeline thus included repeated results in each group - i.e. in an experiment comparing group P, with two scans per individual, to group Q, with three scans per individual, two of the measures for each individual in group Q would be identical to those for the same individual in group P. However, the same experiment involving estimates from the longitudinal processing pipeline had subtly different measures due to the different template depth used to produce the estimates for each group. Thus, group differences detected in measures derived from the cross sectional pipeline could be attributable to the additional measure per individual, while the differences detected in measures derived from the longitudinal pipeline could result from both the additional measure and the change in values resulting from different template depths.

Two classes of bias associated with group difference in template depth were investigated: differences in group mean brain metrics and differences in group trajectory of brain metrics. Differences in group mean were computed from difference in intercepts estimated by regression models without group interaction terms. Differences in trajectories were derived from regression models with group interaction terms (see Section 3.6 for details). These experiments explored whether group differences in metrics could be induced by group differences in template depth.

A second class of analysis explored the potential for systematic bias, namely whether brain metrics or trajectories of brain metrics increase or decrease with template depth. This analysis used a single model fitted to all template depths, rather than pairs of groups as above and provides additional insight by summarizing across all template depths. The simulated atrophy data has constant rates of atrophy with time, and thus brain metrics (such as mean cortical thickness) will be strongly predicted by time. Finding template depth to be an independent predictor of a brain metric in a model including time indicates a likely systematic bias. Bias in trajectory estimates are indicated by a significant interaction between template depth and time.

Both classes of analysis were applied to estimates derived from the cross sectional pipeline and longitudinal pipeline.

A subset of the second class of analysis was applied to real longitudinal ADNI data, rather than simulated atrophy.

### 3.2 MRI data

Two subsets of data from the ADNI1 cohort were selected. *Set A* consisted of participants with MRI acquisitions at the 60 month followup. *Set B* consisted of 50 randomly selected participants from the normal cognition group with baseline scans who were not in Set A. Set B was used as a basis for simulated atrophy. ADNI was launched in 2003 with the primary aim of determining whether serial imaging and other biomarkers could be combined to measure progression of mild cognitive impairment and early Alzheimer’s disease. Written, informed consent was obtained from all participants. Full details of ADNI study design, ethics approvals and acquisition protocols are described at http://www.adni-info.org. This study used the 1.5T MP-RAGE acquisitions from ADNI1 that had been preprocessed using the image correction procedures implemented by ADNI.

### 3.3 FreeSurfer processing of Set A

Six estimates of baseline cortical thickness were obtained for each participant by carrying out six separate runs of the FreeSurfer longitudinal pipeline. Each run used a template with a different depth - i.e. constructed using a different number of acquisitions (ranging from 1 to 6).

Cortical thickness trajectories of Set A are unknown and cannot be assumed to be linear or consistent. Thus analysis of Set A was restricted to the effect of template depth on baseline thickness estimated using the FreeSurfer longitudinal pipeline.

FreeSurfer v6.0 was used for all experiments without manual editing.

### 3.4 Simulated MRI data

Scans from Set B were used as input to the Simulatrophy tool (Khanal et al. 2016; Khanal, Ayache, and Pennec 2016) (available from https://inria-asclepios.github.io/simul-atrophy/) to create longitudinal data with simulated atrophy of cortical gray matter.

Five time series, each containing six time-points, were created for each baseline scan. A different rate of atrophy, ranging from 0 to 4% (settings 00, 01, 02, 03 and 04) was used for each time series. A cortical mask derived from the FreeSurfer parcellation of the original baseline scan was used to define brain regions experiencing atrophy.

The simulated data was thus constructed to have a known and constant rate of atrophy. All simulation data was derived from the baseline scan of each participant and was thus of similar quality to the baseline scan.

To determine the effect of simulating WM atrophy on cortical thickness estimates, we repeated our analysis with atrophy restricted to WM (See Supplementary A.1.2). The region experiencing atrophy was defined by a 1 voxel erosion of the white matter mask derived from FreeSurfer.

### 3.5 FreeSurfer processing of simulated data

Six longitudinal FreeSurfer analyses, each with a different template depth, were carried out for each time series. All of the acquisitions contributing to each template were processed longitudinally, rather than only the baseline scan, as was done with Set A.

Longitudinal FreeSurfer processing begins with cross sectional processing of all acquisitions and thus standard, cross-sectional, estimates of thickness were available as a result of this procedure. The mean cortical thickness and supratentorial volume estimates were used in the analysis.

### 3.6 Statistical Analysis

Multi-level models, with random intercepts per participant to account for individual difference from the group average, were used to analyse all experiments due to the repeated measure design. Model formulae, using R syntax, are in Supplementary Section A.2.1. Random intercept models were used for all simulated atrophy experiments as the atrophy was held constant (i.e. identical for different individuals), making random slope models unnecessary.

Trajectories of cortical thickness and supratentorial volume estimated with cross sectional processing for scans with simulated atrophy were analysed separately for each atrophy level and type. These models provide a group estimate of slope to validate the performance of simulation.

Effects of template depth on the longitudinal pipeline were investigated by comparing mean thickness and trajectories (change of measures with time derived from regression models) estimated using templates of different depth. Using template depth as a group main effect, we fitted models predicting thickness from time point, for example the trajectories estimated from a template depth of 3 were compared to those estimated from template depth 6. All possible template depth pairs (10) were explored for each atrophy type and level.

The equivalent models based on cross sectional FreeSurfer estimates were fitted for comparison purposes. No correction for multiple comparisons was performed for the depth pair analysis as the intention was to explore the range of scenarios that may occur in practice. A p-value of below 0.05 was taken to indicate a group difference.

Template depth was also used as a continuous main effect and continuous interaction term to investigate whether it was an independent predictor of mean thickness or trajectory and thus test for systematic bias.

A multi-level model was used to test whether the estimate of baseline thickness from the longitudinal pipeline applied to Set A was associated with template depth.

Cohen’s f was used to characterize effect size.

Analysis was carried out in R version 4.0 (R Core Team 2020) and the lme4 package (Bates et al. 2015). Model formulae are provided in Supplementary Material Section A.2

## 4 Results

### 4.1 Demographics

Scans included in Set A were acquired from 56 participants (19 female) with followup scans at 60 months. Most participants (36) had 6 scans, 13 had 5, 5 had 4 and 2 had 3 scans. Mean age at baseline was 75.1 (sd=5.4), 31 were cognitively normal at baseline and 25 had a diagnosis of mild cognitive impairment.

Simulated scans in Set B were generated from a in independent random sample of 50 (26 female) cognitively normal ADNI participants with a mean age of 76 at baseline. (sd=4.9).

### 4.2 Impact of template depth on cortical thickness estimates in ADNI

#### 4.2.1 Set A: ADNI acquisitions to 60 months

Cortical thickness trajectories for Set A estimated using the cross sectional and longitudinal FreeSurfer pipelines are shown in Figures 1 and 2. A range of rates of atrophy are apparent in the sample and trajectories are noisy, as expected in a real longitudinal study. The longitudinal pipeline results in Figure 2 illustrate that there is variability resulting from differences in template depth as thickness estimates at the same time point do not overlap perfectly, however no clear pattern of difference is discernible.

**Figure 1:**
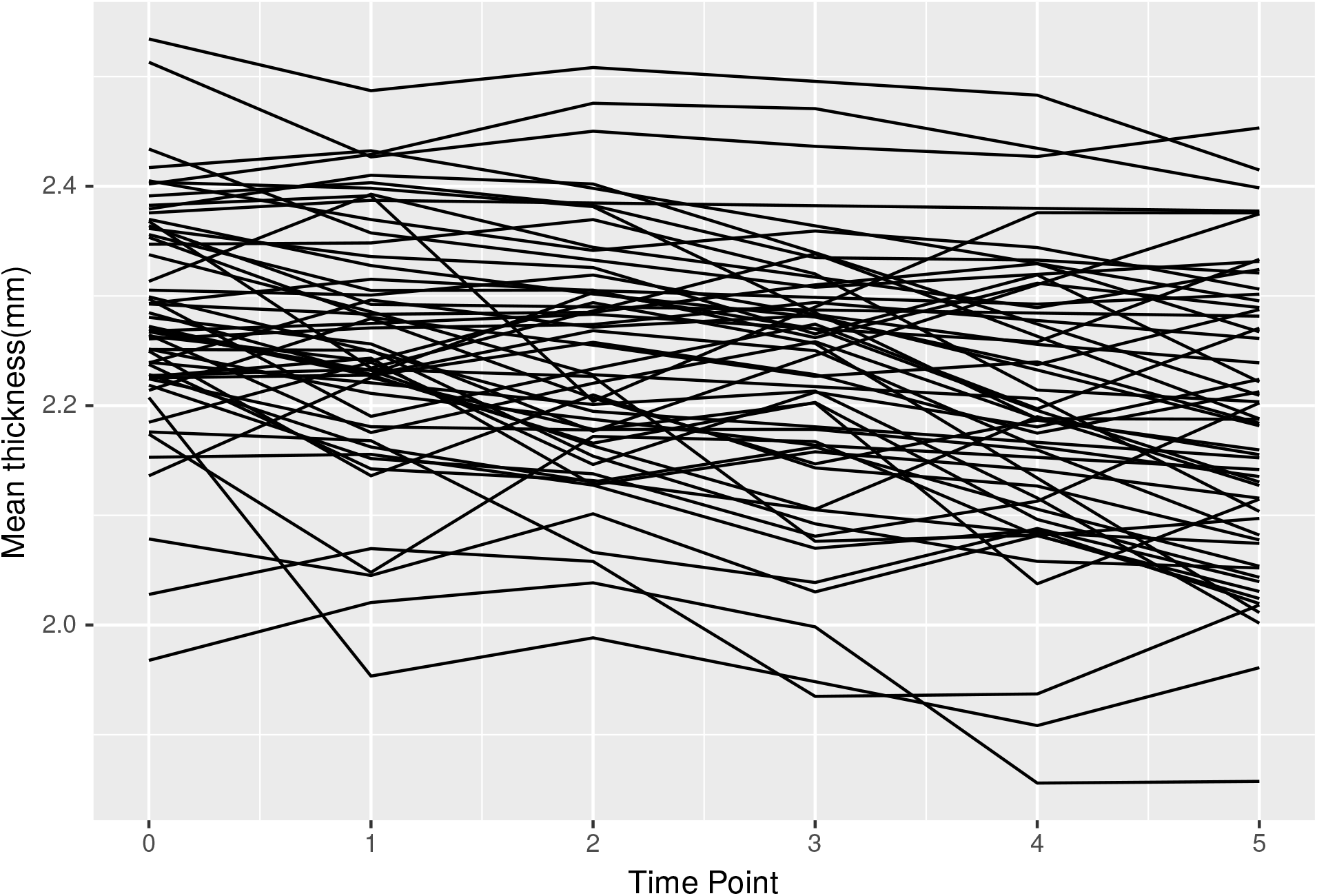
Mean cortical thickness estimated with cross sectional FreeSurfer for ADNI participants with scans at 60 months.

**Figure 2:**
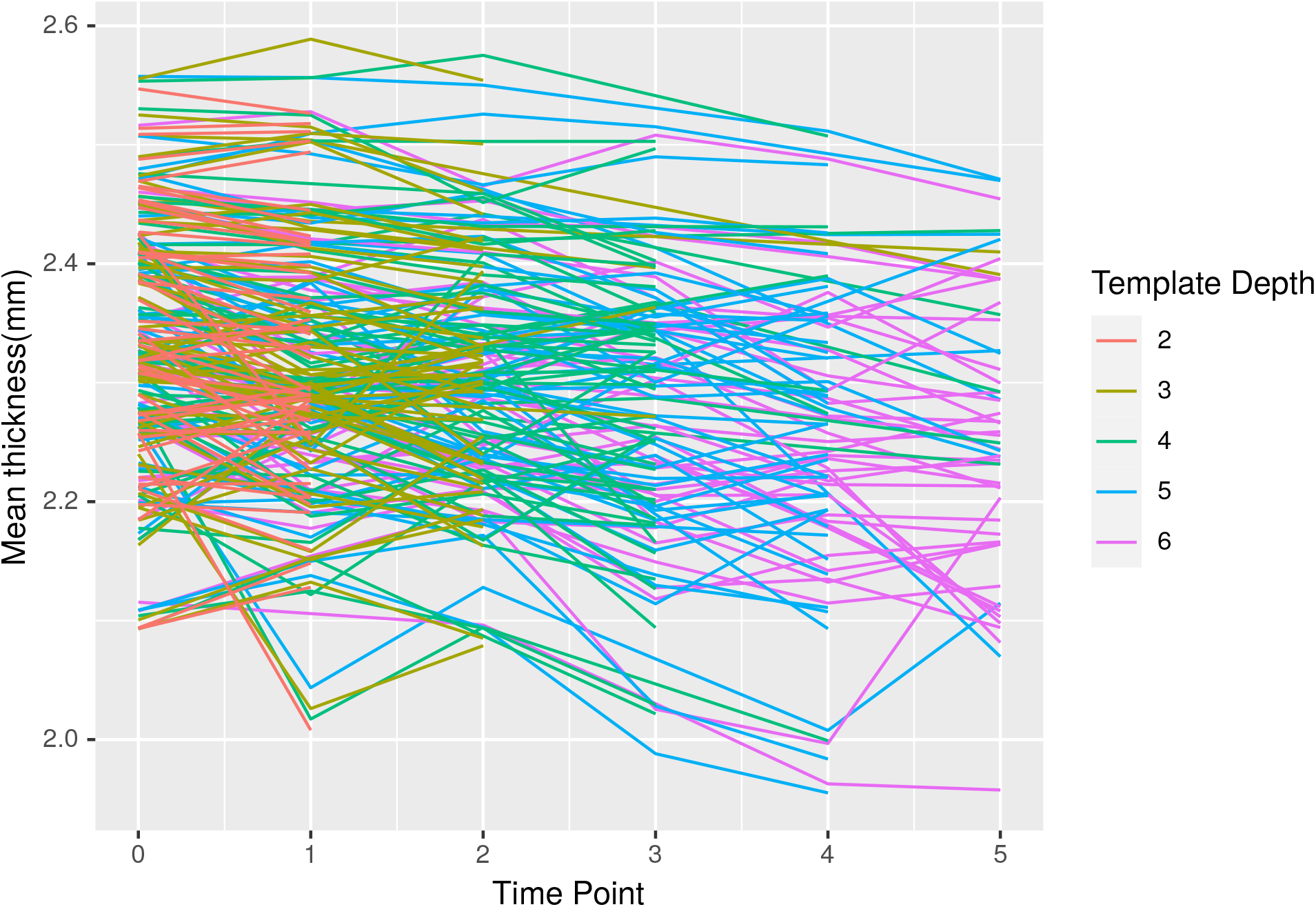
Mean cortical thickness estimates with longitudinal FreeSurfer and various template depths for ADNI participants with scans at 60 months. Each line represents a set of cortical thickness estimates for a given subject, coloured by the number of MRI scans used in the longitudinal pipeline to generate that estimate. The variation of mean thickness estimates caused by different template depths is visible.

The baseline thickness estimates produced by the longitudinal pipeline with different template depths and the estimated baseline thickness as a function of template depth are shown in Figure 3. We tested whether the template depth had a significant effect on baseline cortical thickness estimates produced by the longitudinal pipeline. We found that template depth was a significant predictor of mean cortical thickness at baseline in Set A (p=5.9e-11), coefficient 0.0026 (stderr 0.0004) and a Cohen’s f statistic of 0.43. We found that template depth was not a predictor of supratentorial volume (p=0.717).

**Figure 3:**
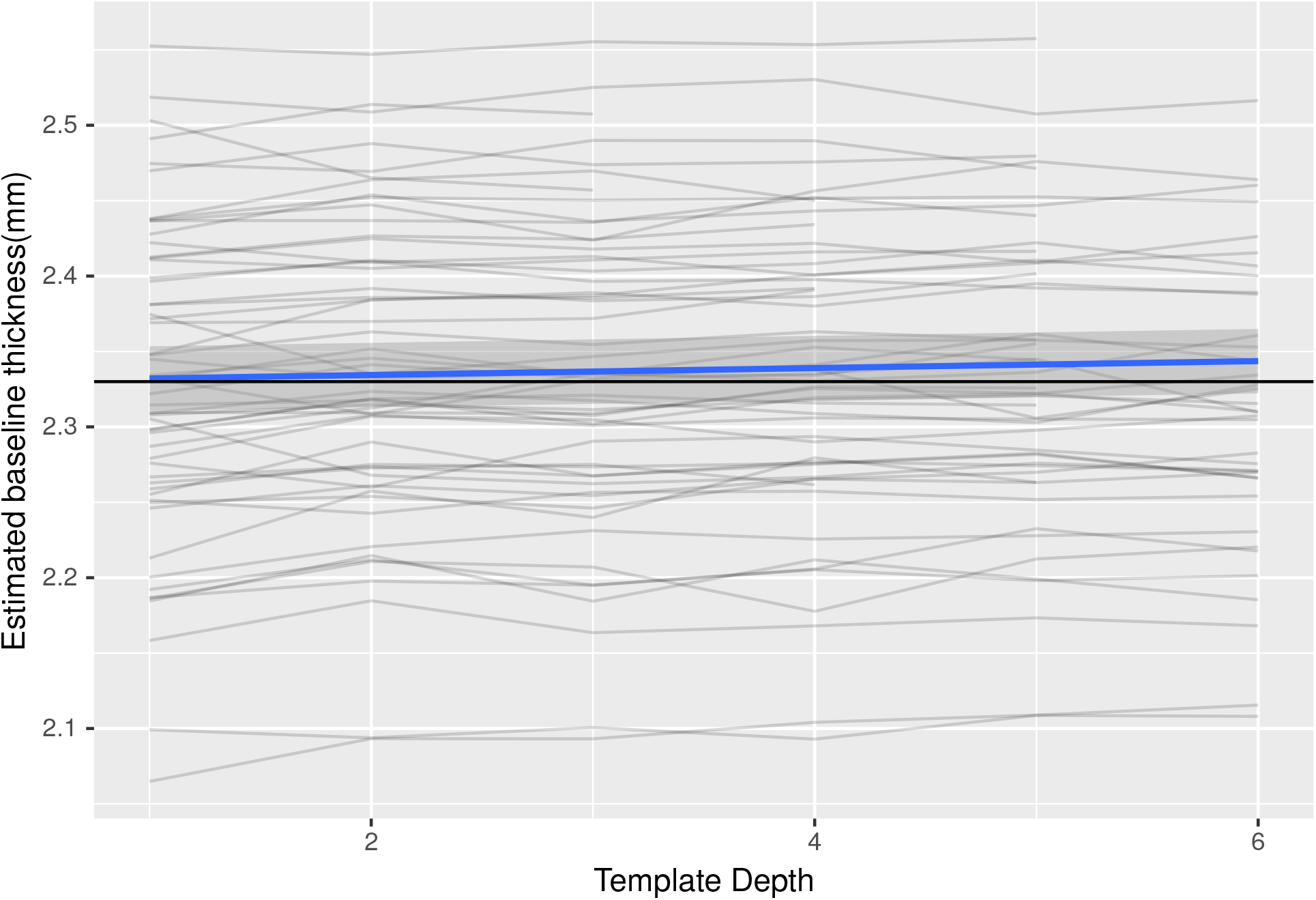
Mean cortical thickness at baseline estimated with longitudinal FreeSurfer and different template depths for ADNI participants with scans at 60 months. Each line represents estimates of baseline cortical thickness for a given subject, based on longitudinal processing of data with increasing template depths. With no bias, estimates should be stable within individuals across template depths. The blue line illustrates the global estimate of change in thickness estimates for baseline scan as a function of template depth (*β* = 0.0026mm per additional image in the template). The black line is for comparison purposes and is horizontal.

### 4.3 Set B - Simulated atrophy of randomly selected, cognitively normal, ADNI participants

#### 4.3.1 Simulated cortical atrophy

We simulated linear, global cortical atrophy in a sample of 50 cognitively normal ADNI particpants. Trajectories of cortical thickness and supratentorial volume produced by cross sectional FreeSurfer processing of each atrophied scan are shown in Figure 4 and Supplementary Figure S2. Each panel illustrates a different atrophy setting. Non-zero atrophy settings induce decreases in cortical thickness estimates across all simulated examples and detected changes are proportional to atrophy settings. On average, cortical thickness decreased by 0mm, 0.0033mm, 0.008mm, 0.011mm, 0.015mm per timepoint (p < 0.0001 for non-zero atrophy settings) for atrophy settings 0,1,2,3,4 respectively. The range of atrophy levels is broadly similar to that observed in the real data while the individual trajectories are less noisy, as expected of simulated atrophy.

**Figure 4:**
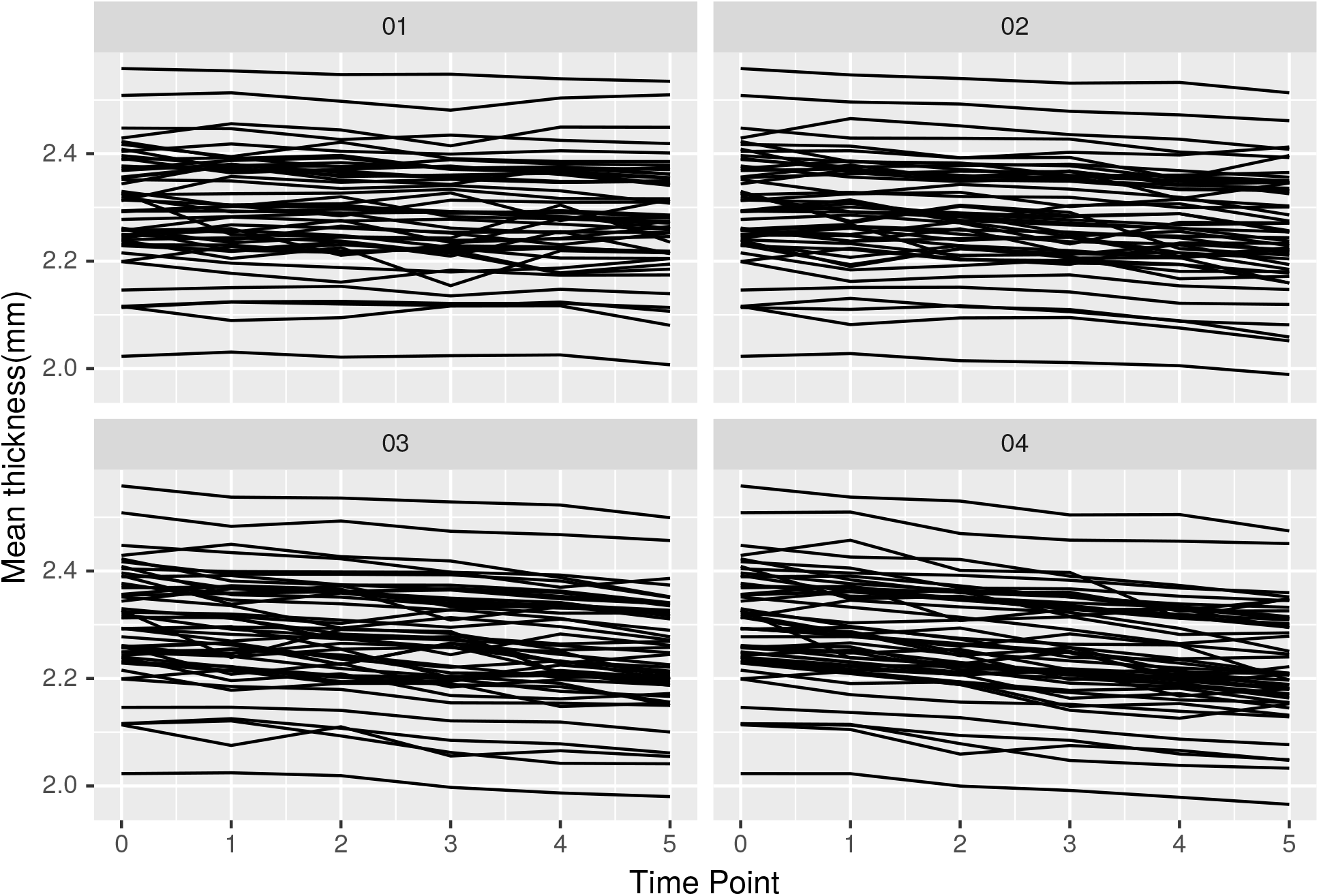
Mean cortical thickness estimated with cross sectional FreeSurfer in presence of simulated cortical atrophy. Each panel corresponds to a different level of simulated atrophy.

An example of the surfaces extracted from two time points for the same participant with simulated cortical atrophy is illustrated in Figure 5. This example illustrates time points 1 and 5 and an atrophy level of 04. These figures demonstrate the subtle change caused by high atrophy settings and the difficulty interpreting change using visualization, especially at a local level in a single slice. At higher levels of simulated cortical atrophy there is a disparity between both the WM/GM and GM/CSF boundaries. In some cases the red surface is outside the blue surface, despite atrophy leading to measurable global cortical thinning. This effect is caused by slight movement of gyrus as a result of simulated atrophy.

**Figure 5:**
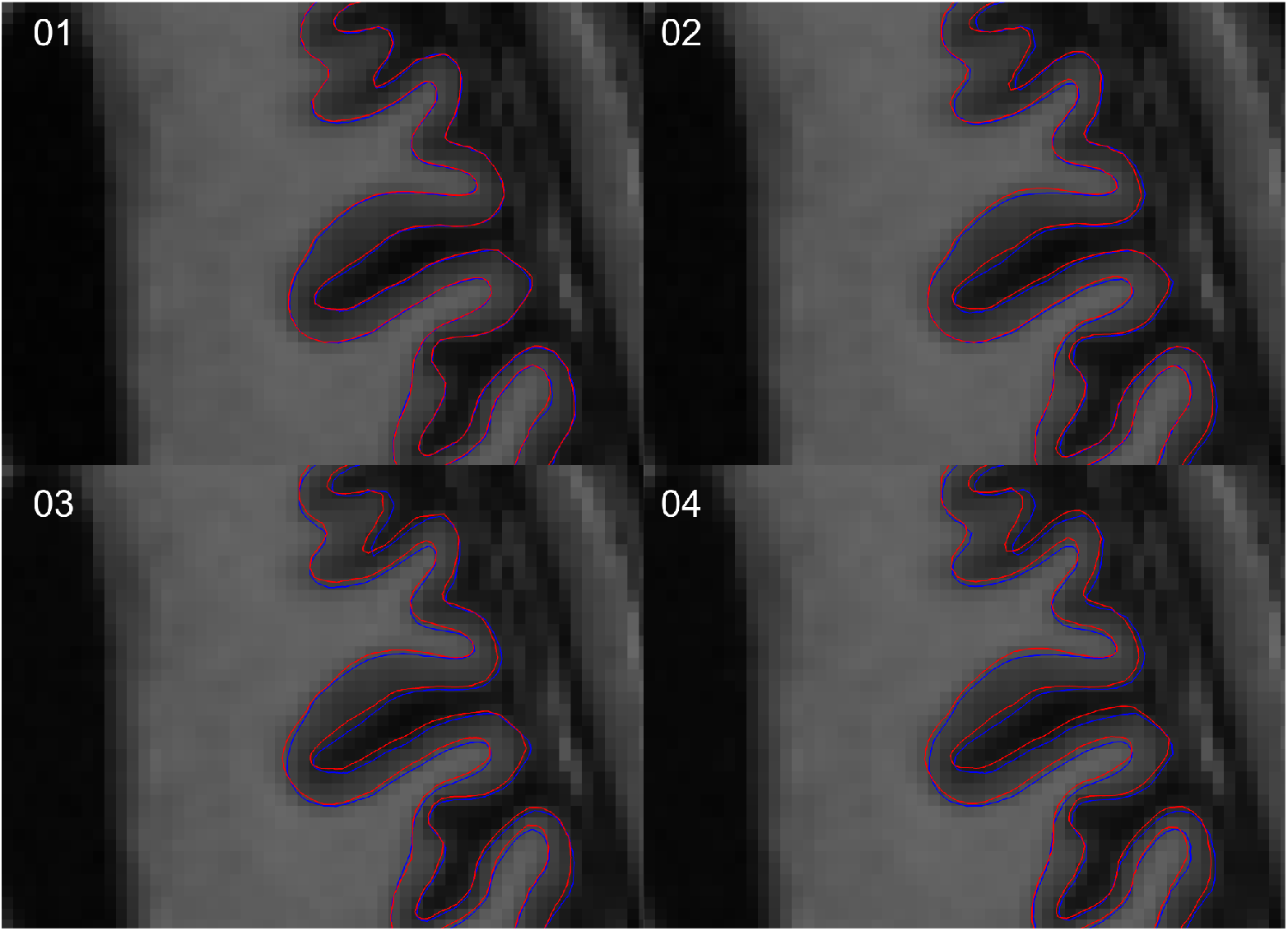
FreeSurfer WM and pial surfaces corresponding to simulated cortical atrophy time points 1 (blue) and 5 (red) for atrophy levels 01 to 04. These figures illustrate the difficulty in visually interpreting atrophy at a local level. Apparent switching of boundaries (red outside blue) occurs as a result of gyral shift in the simulation process.

### 4.4 Variation in simulated cortical thickness estimates using the longitudinal pipeline

#### 4.4.1 Simulated cortical atrophy

The simulated trajectories in Figure 4 were recalculated using the longitudinal FS pipeline, using different template depths.

Trajectories of mean cortical thickness and supratentorial volume produced by longitudinal FreeSurfer processing in the presence of simulated cortical atrophy are shown in Figure 6 and Supplementary Figure S5. The trajectories for an individual participant differ with template depth, but group level patterns are not evident from visualization alone.

**Figure 6:**
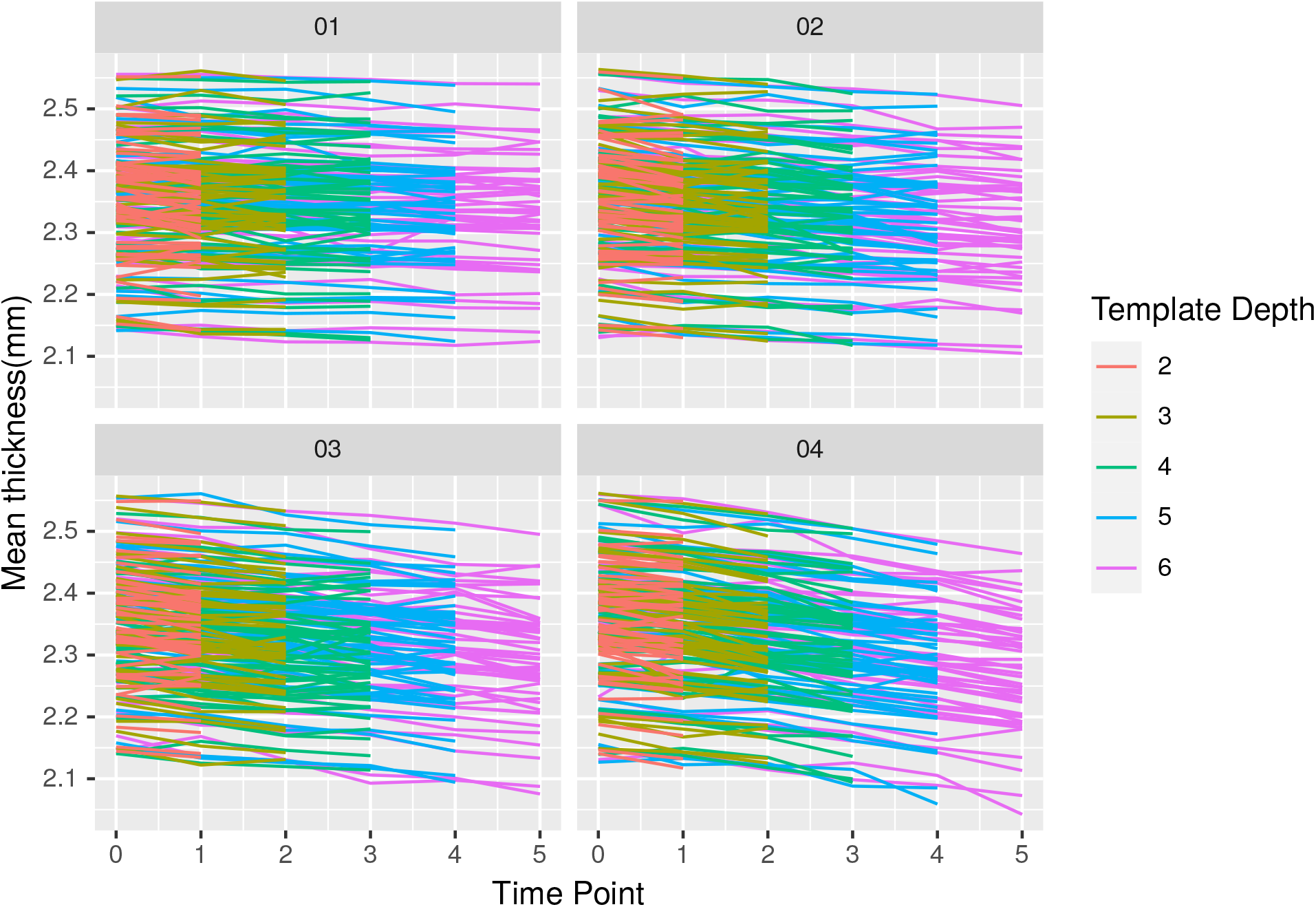
Cortical thickness estimated with longitudinal pipeline and varying numbers in template in presence of simulated cortical atrophy. Each panel corresponds to a different level of simulated atrophy. Each line represents cortical thickness trajectories for a given simulated subject, coloured by the number of MRI scans used to estimate thickness in the longitudinal FreeSurfer pipeline.

### 4.5 Comparison of template depth groupings

Group comparisons are presented in tabular form in Figures 7 to 10 and S8 to S11. Each figure panel corresponds to a different atrophy level. Each cell represents a comparison of mean cortical thickness or thickness trajectory estimated in the simulated data using groups of the same individuals with different template depths. This comparison was based on the group main effect or interaction term in the multilevel models. Positive beta values indicate that the coefficient for the group with a higher template depth was larger than the coefficient for the group with the lower template depth. As the simulated data for a given degree of atrophy underlying each comparison is the same, we would expect no significant differences if the longitudinal pipeline provides consistent estimates of baseline thickness/trajectory for each template depth. Tests comparing images processed using the cross sectional pipeline (i.e.: each image processed separately without reference to an individualised template) are in the upper left of each panel while the lower right contains results of tests using data from the longitudinal pipeline. Each cell contains the p-value, beta coefficient and Cohen’s f statistic resulting from a single group comparison. Red text indicates p-value below 0.05 and bold and italic text indicate negative and positive beta coefficients respectively.

**Figure 7:**
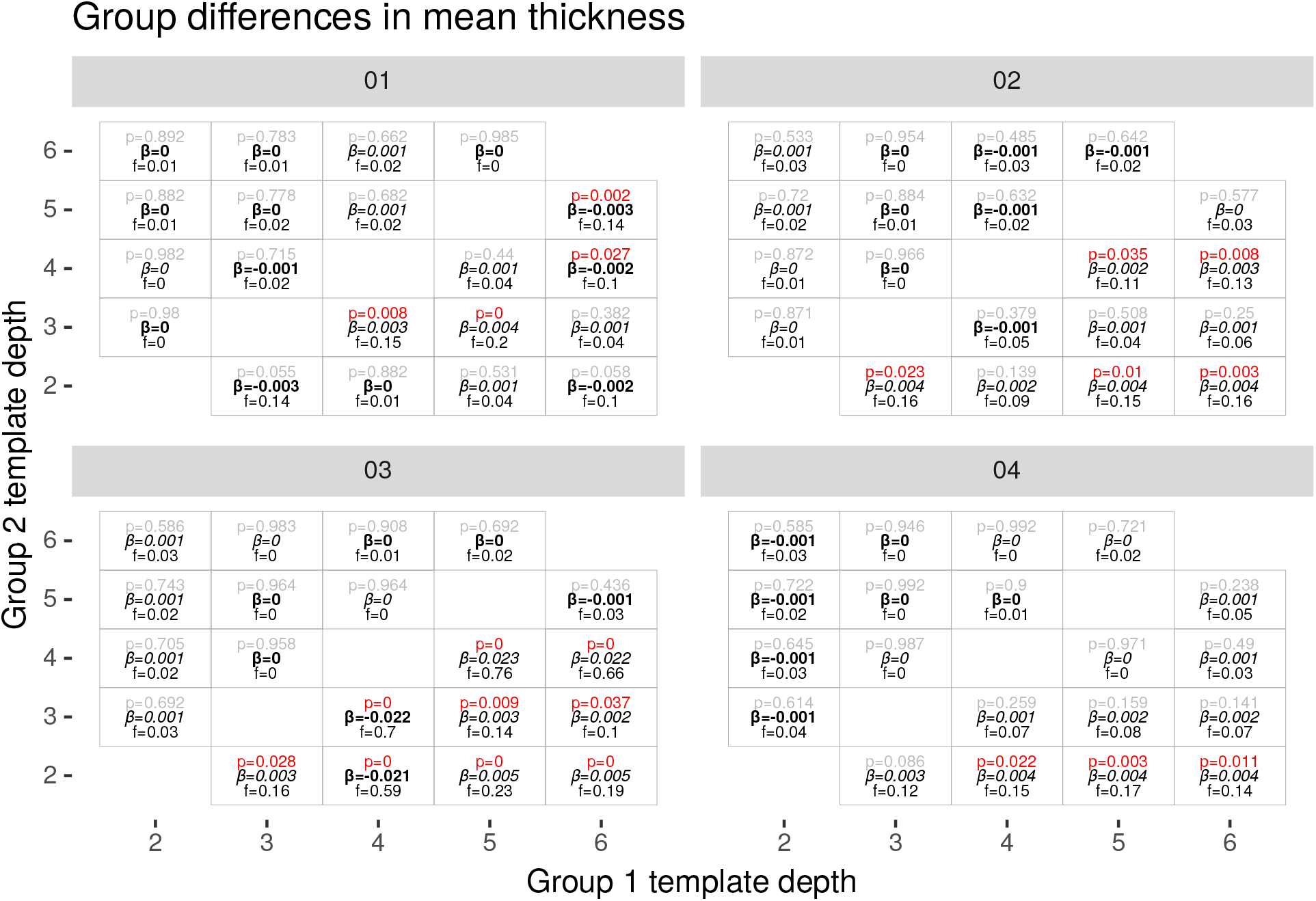
Group differences in cortical thickness for simulated GM atrophy. Positive *β* indicates a higher coefficient in the group with higher template depth.

**Figure 8:**
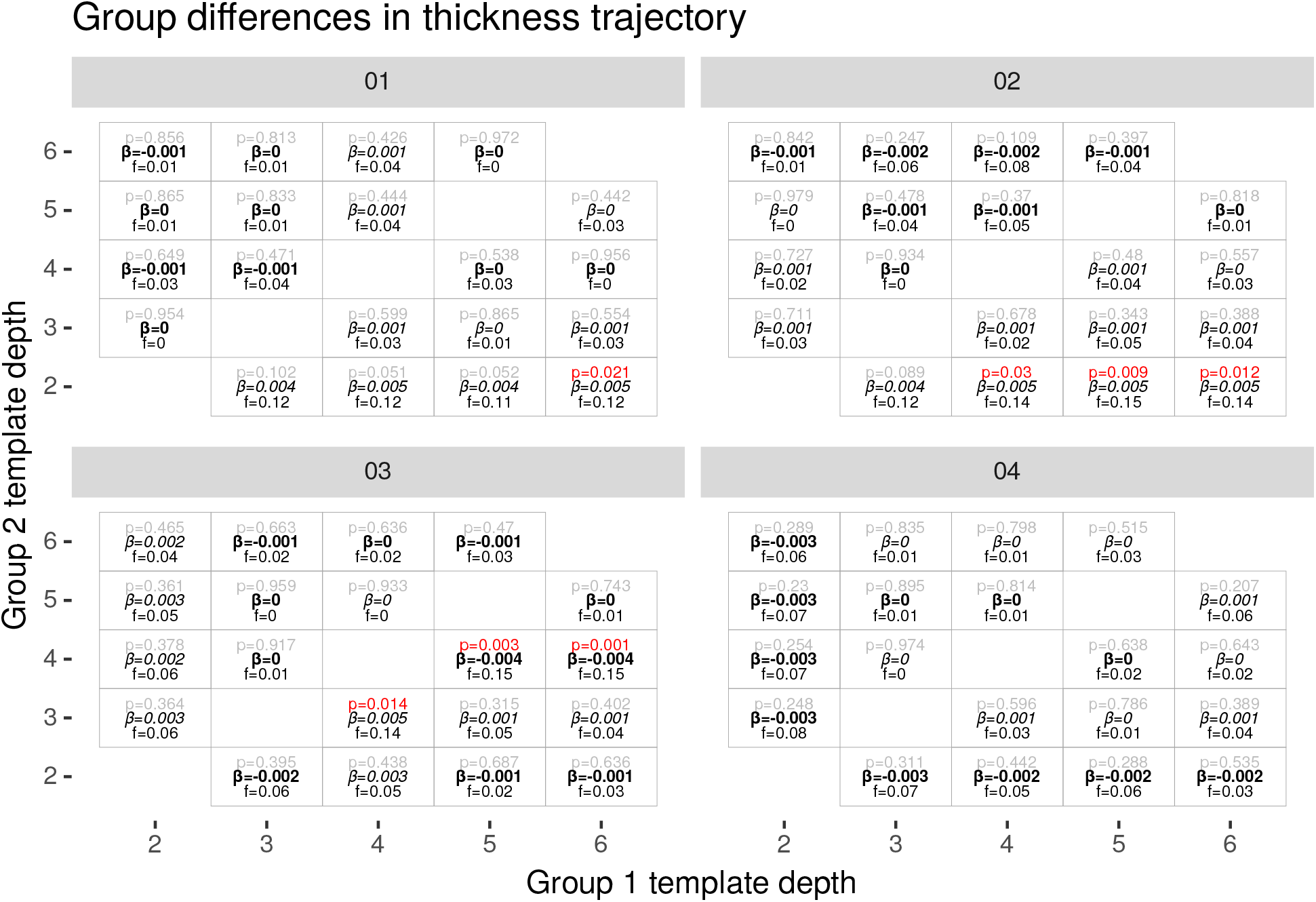
Group differences in trajectory of cortical thickness for simulated GM atrophy. Positive *β* indicates a higher coefficient in the group with higher template depth.

**Figure 9:**
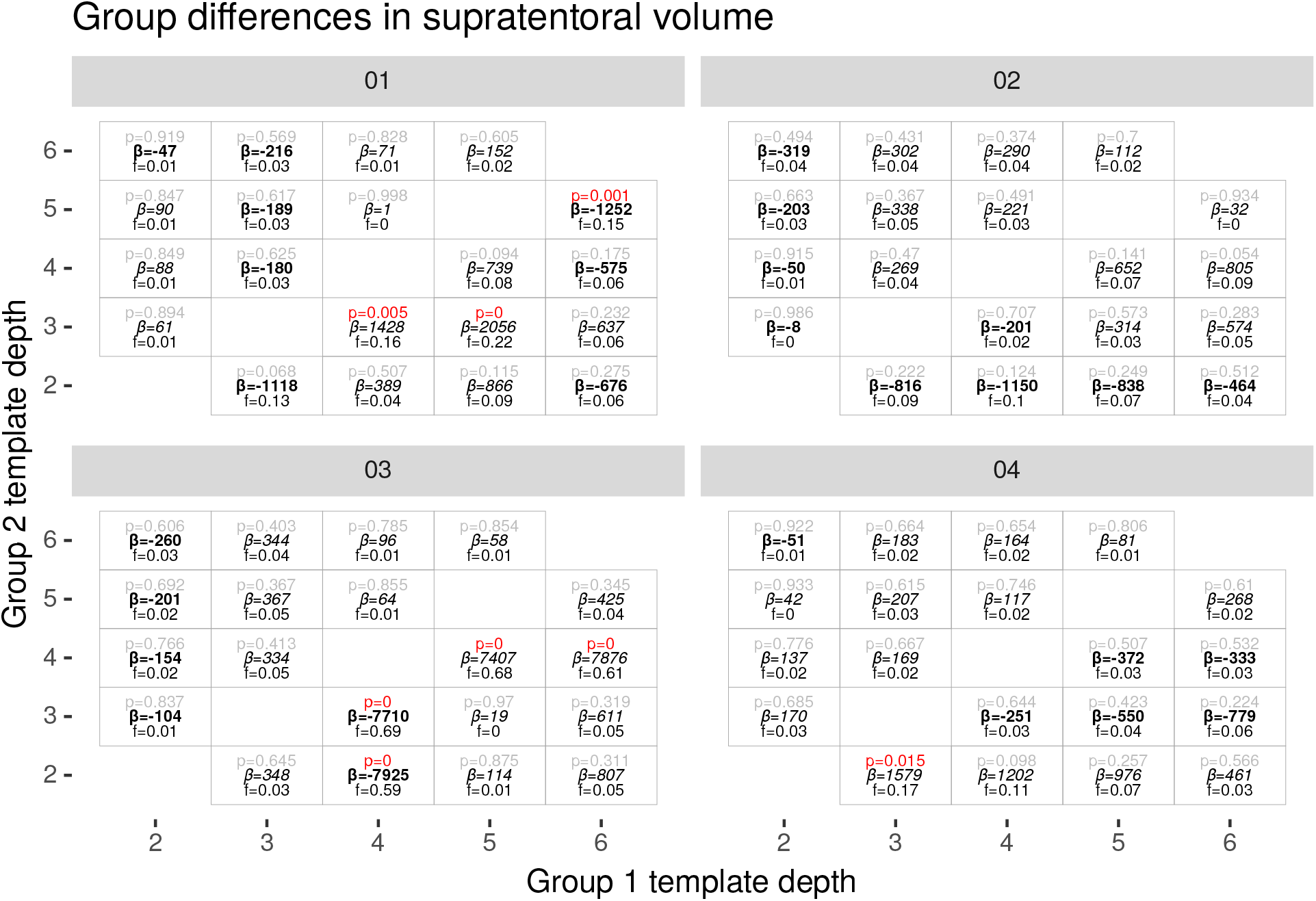
Group differences in supratentorial volume for simulated GM atrophy. Positive *β* indicates a higher coefficient in the group with higher template depth.

**Figure 10:**
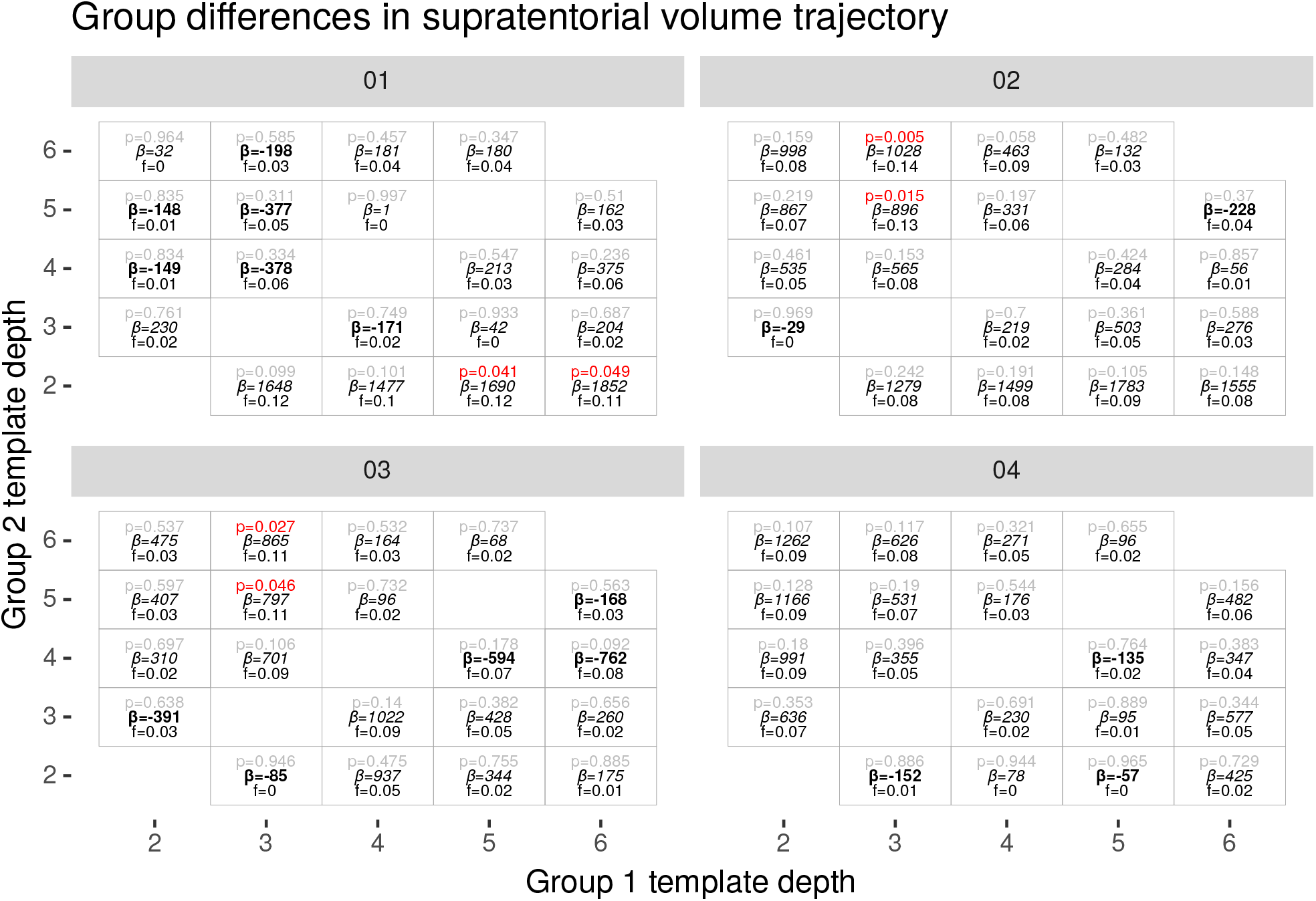
Group differences in trajectory of supratentorial volume for simulated GM atrophy. Positive *β* indicates a higher coefficient in the group with higher template depth.

#### 4.5.1 Cortical thickness

No template depth group differences were detected in mean GM thickness or trajectory (minimum p value 0.07) for either cortical atrophy or WM atrophy when individual images were processed using the cross sectional FreeSurfer pipeline

Differences in mean cortical thickness between template depth groups were detected (p < 0.05) in 26 of the 50 comparisons (10 group pairs, 5 levels of atrophy) of simulated cortical atrophy processed using the longitudinal pipeline (Figure 7).

The minimum p-value (p=1e-41) occurred when comparing template depths 4 and 5 at atrophy level 03 and corresponded to a difference in means of 0.023mm (stderr 0.00149) in 2.33 (Cohen’s f = 0.76).

The sign of group differences was not consistent, i.e. more acquisitions did not imply thicker (or thinner estimates)

Differences in trajectories of cortical thickness between template groups were observed in 7 of the 50 comparisons (Figure 8).

The minimum p-value (p=0.0012) occurred when comparing template depths 4 and 6 at atrophy level 03. Group difference in trajectory = -0.00374mm/TP and the slope of base group = -0.008 (Cohen’s f = 0.15). (almost 50% difference)

The sign of group differences in trajectory of cortical thickness was not consistent.

Similar results were observed when applying atrophy to WM only - See Supplementary Material Section A.3.

#### 4.5.2 Supratentorial volume

In contrast to the cortical thickness results, several significant differences in trajectory of supratentorial volume estimated from cross sectional processing were detected: 3 vs 5 time points (p=0.015) and 3 vs 6 time points (p=0.005) for cortical atrophy level 02, and 3 vs 5 time points (p=0.046) and 3 vs 6 time points (p=0.027) for cortical atrophy level 03. See upper left triangle of Figure 10.

Differences in mean supratentorial volume between template depth groups were detected (p < 0.05) in 8 of the 50 comparisons of of simulated cortical atrophy processed the longitudinal pipeline (Figure 9).

The minimum p-value (p=2e-34) occurred when comparing template depths 4 and 5 at atrophy level 03. The group difference in mean was 7407(stderr 549) in 898576 (Cohen’s f = 0.68).

The sign of group differences was not consistent.

Marginally significant differences in trajectory of supratentorial volume were detected in 2 of 50 comparions of simulated cortical atrophy (Figure 10). The minimum p-value (p = 0.041) occurred when comparing template depths 2 and 5 at atrophy level 01. The group difference was 1690(stderr 825) and the slope of base group = -2785. (Cohen’s f = 0.12)

Sign of group differences was positive for both cases.

### 4.6 Template depth as a predictor of brain metrics

#### 4.6.1 Set A

Template depth was a significant predictor of mean cortical thickness in Set A (p=1e-9), coefficient 0.0026(stderr 0.0004) and a Cohen’s f statistic of 0.42. See Figure 3.

#### 4.6.2 Set B

##### 4.6.2.1 Cortical atrophy

We modeled mean cortical thickness as a function of template depth using multi-level models. Template depth was a significant predictor of mean cortical thickness at atrophy levels of 02 (*β* = 0.0009, p=0.002, Cohen’s f = 0.1), 03(*β* = 0.002, p<0.00001, Cohen’s f = 0.15) and 04(*β* = 0.001, p=0.002, Cohen’s f = 0.10) and of supratentorial volume for atrophy level 03 (*β* = 545, p=0.002, Cohen’s f = 0.10). It was also a predictor of cortical thickness trajectory (*β* = 0.001, p=0.004, Cohen’s f = 0.09) and supratentorial volume trajectory (*β* = 536, p=0.0001, Cohen’s f = 0.13) at an atrophy level of 03. Coefficients were positive - deeper template corresponding to thicker cortex or larger volume.

##### 4.6.2.2 WM atrophy

Template depth was a predictor of mean cortical thickness and supratentorial volume for atrophy levels 02, 03 and 04 (*β* =−0.001, −0.002, −0.002 and *β* =−572, −688, −773, p < 0.0001, Cohen’s f = 0.14, 0.22, 0.17 and f = 0.13, 0.15, 0.14). Coefficients were negative.

Template depth was positively associated with cortical thickness trajectory for atrophy levels 00 and 04 (*β* = 0.0004, 0.0005, p=0.01, 0.03 and Cohens f=0.08, 0.07).

## 5 Discussion

In this study we have investigated the impact of *template depth*, the number acquisitions used to construct individual templates, on measures produced by the FreeSurfer longitudinal pipeline. It is rare for longitudinal neuroimaging studies to have a balanced design, with the same number of acquisitions for every participant. Participants may drop out of studies for many reasons, some of which may be directly or indirectly related to study questions or exposures. This is particularly common in studies of aging or chronic/progressive disease that may limit participants’ ability to complete demanding study protocols, leading to lower rates of follow-up in the sicker or frailer portions of the study cohort. Other biases in followup-rates caused by, for example socio-economic-status, have the potential to affect studies of typical development or healthy controls. One can imagine diverse potential causes - for example a group-dependent frequency of dental braces could lead to a group difference in number of usable scans acquired during adolescence while differing rates of employment may lead to increased or decreased attendance, depending on study type. Difference in followup rates of study groups will lead to group differences in template depth. It is therefore important to understand the effect of template depth on derived measures.

This article has investigated the problem via two sets of data derived from the ADNI study. We created several simulated longitudinal datasets derived from real scans using a previously described atrophy simulator (Set B). The datasets were constructed using atrophy rates that remained constant for the course of the simulation (i.e constant over time) and with the same rate and pattern for each individual. We investigated two patterns of atrophy, one cortical specific and one white matter specific. The visualization of cross sectional processing results suggests that Simulatrophy, with a cortical atrophy pattern, produced rates of cortical thinning that remained constant over the 6 simulated time points and were linearly related to the atrophy setting. The WM atrophy pattern did not produce detectable cortical thinning in cross sectional processing results at lower atrophy levels, but did begin to influence cortical thickness measures in a detectable manner at atrophy settings of 03 and 04. This is likely to be due to the spatial regularization required to produce realistic physical models and partial volume effects at the GM-CSF boundary. The highest rate of cortical thinning detected with the WM atrophy pattern was slightly lower than that detected with the lowest nonzero cortical atrophy pattern.

These simulated longitudinal acquisitions represent an ideal data set for FreeSurfer processing, as the non brain tissue was not modified by the atrophy simulator and the contrast to noise and bias field inhomogeneity effects remained constant within a participant.

Significant differences in mean cortical thickness and trajectories between template-depth groups were more frequently detected in the results of the FreeSurfer longitudinal pipeline than the standard cross-sectional pipeline. The differences that were detected in the cross sectional estimates were marginally significant and involved supratentorial volume rather than cortical thickness. As each scan is processed independently by the cross sectional pipeline, any differences detected using estimates from the cross-sectional pipeline must result from noise. The lack of differences we detected indicate a low rate of type 1 errors (false positives), providing a reference for the frequency of differences detected using estimates from the longitudinal pipeline.

Differences in mean thickness were detected more frequently than differences in thickness trajectory. Signs of mean thickness difference appear to depend on the type of atrophy, with deeper templates associated with lower thickness in the white matter atrophy simulation but no consistent relationship present in the cortical atrophy simulation. Differences in trajectory appear to be less common, fortunately, as the effect of template depth on trajectory appears weaker than on mean thickness. However, the results in Section 4.6 indicate that the effect is detectable under some conditions and is positive. The effect sizes of group differences in trajectory in the presence of GM atrophy were small and the signs were inconsistent. One marginally significant difference in thickness trajectory was detected in the white matter atrophy simulation, occurring at an atrophy level at which the thickness trajectory was zero in the cross sectional data.

These results are complex. It is clear that group differences in mean cortical thickness, trajectory of cortical thickness, mean supratentorial volume and trajectory of supratentorial volume can be induced via the choice of template depth used in longitudinal image processing. Differences in mean thickness appear to be more common than differences in trajectory, and occurred at similar rates in the presence of either cortical or white matter atrophy, supported by our finding that template depth was a significant predictor of mean thickness in both atrophy experiments. Supratentorial volume measures exhibited a similar behaviour.

We have examined the effect of template depth using both group comparisons (Section 4.5) and as a predictor of brain measures (Section 4.6). The coefficients in Section 4.6 were highly significant in some cases, with Cohen’s effect sizes in the small-medium range. However the magnitude of the coefficients were small, corresponding to 0.1% increase in thickness per unit of template depth in Set A and similar or smaller values in Set B. These values were estimated using all combinations of available data and indicate that differences attributable to template depth are not random - i.e. bias is present.

The small magnitude of the bias may appear to be of minimal interest to practical studies. However the group-wise comparisons indicate that there is cause for concern. The largest difference in cortical thickness between groups resulting from differences in template depth was 0.023mm, or approximately 1%. This is a potentially large confound in studies of for example, of typical development where the change in cortical thickness between ages 5 and 25 is approximately 9% (Mills and Tamnes 2014) and differences between groups over this period will typically be smaller, or diseases like type 2 diabetes mellitus, where effects on cortical thickness may be of similar magnitude to the template depth induced difference (0.03mm)(Moran et al. 2015).

Accuracy and test-retest reliability of the FreeSurfer, for the purposes of sample size calculations, have been investigated previously (Liem et al. 2015; Pardoe et al. 2013). These articles examined the problem in the context of a vertex-wise analysis with varying levels of smoothing and a mix of acquisition models, with (Liem et al. 2015) using repeated scans and the longitudinal FreeSurfer pipeline while (Pardoe et al. 2013) cross sectional studies and group comparisons. The analysis by (Liem et al. 2015) suggests that a difference of 0.053mm can be detected between groups 20 if a large smoothing kernel (20mm) is used. The power calculation tool they provide suggests that a 0.023mm difference is detectable in a pairwise vertex based analysis with N=76 and 20mm smoothing. Studies examining global mean thickness, i.e. a very large smoothing, like ours, and more than two timepoints can expect a higher sensitivity. The magnitude of thickness differences resulting from differences in template depth is thus large enough to be of potential concern in some studies.

The effect of template depth on cortical thickness estimates was also observed in real longitudinal data from the ADNI study. Repeated analysis of baseline scans using templates of different depth, constructed from real (not simulated) follow up scans, demonstrated that template depth was a predictor of baseline cortical thickness, and that the effect size was large (f=0.42).

The Cohen’s f effect sizes for significant group differences were typically in the range 0.1-0.2 (small) for cortical thickness in the presence of GM atrophy, while moderate effects (over 0.25) were observed in the presence of WM atrophy. Occasional large effect sizes (>0.6) were observed for tests of supratentorial volume in the presence of GM atrophy. Effect sizes for prediction of thickness or supratentorial volume from template depth were typically small.

Trajectories are often the property of interest in longitudinal studies, and fortunately our study suggests that the impact of difference in template depth on trajectories of thickness is not as strong or consistent as the impact on mean thickness. However we have demonstrated that detectable differences can be induced. Group differences in mean thickness, while typically not the main aim of longitudinal studies, are frequently reported and are an important component of group characteristics. We have detected an association between template depth and mean thickness in both real data and a simulated white matter atrophy data set.

We can speculate on possible underlying causes of the bias we have detected. The tendency of templates created via iterative registration procedures to be larger than the average size of the contributing images is known as *template drift* and is well documented (Abdollahi et al. 2014; Van Essen et al. 2011; Buckner et al. 2004). The individual templates used by the FreeSurfer longitudinal pipeline are created via an iterative process, and may be susceptible to template drift. The relationship between template drift and the bias we have detected is currently under investigation. Another possibility is that the small number of samples involved in creating a template leads to discretization effects, where adding a single sample may lead to a sudden change in template appearance and thus pipeline behaviour.

The FreeSurfer longitudinal pipeline is very effective at improving segmentation reproducibility (Jovicich et al. 2013) and maximizing the information that can be extracted from any one scan based on information from other acquisitions of the same individual at different study timepoints. Great care has been taken to ensure that the templates used are not biased by one acquisition. The potential for bias that we have discussed is real and no technical solution is available to eliminate it. The best current advice is to carefully monitor group differences in distribution of template depth, and relationships between template depth and study exposures, in the same way that other, non imaging, study measures must be carefully monitored for potential sources of bias. Magnitude of any effects detected must be scrutinized to determine whether they may be attributable to group differences in template depth, rather than biological or treatment effects. The option of statistically adjusting for template depth may be valid in some studies, but will need to be considered carefully due to correlation with other study variables, such as age. Another option may be to use a fixed template depth, for example in a study with a mix of 3 and 4 scans per individual an attractive option could be to use the first 3 scans to construct the individual template, gaining most of the improved robustness. However the current FreeSurfer implementation requires that scans must be included in the template in order to use the longitudinal pipeline.

### Limitations

This is largely a simulation study with longitudinal data derived from a set of real baseline scans. Simulation is the simplest way to provide known patterns and rates of change. However the limitation is that the variation between acquisitions is lower than might be expected with real data, with scanner drift and changes in noise characteristics, orientation, bias field inhomogeneity and non brain tissue. The FreeSurfer longitudinal pipeline improves segmentation reliability in the presence of such inter-acquisition differences. The absence of such effects in our simulations may have strengthened the signal due to template depth compared to what may be expected in a real study. However, we were able to detect the association between mean thickness and template depth in a real study.

## 6 Conclusion

There is an association between template depth and estimates of mean cortical thickness and supratentorial volume produced by the FreeSurfer longitudinal pipeline. This study has detected the association in both real and simulated data sets. A less consistent association was detected between template depth and cortical thickness trajectories, and we were able to detect differences in trajectory between groups that differed by template depth.

These findings illustrate yet another factor, distinct from established issues relating to selection bias, that must be carefully considered when interpreting the results of longitudinal studies, as differences in followup rates can have direct impact on the results of processing pipelines.

## 7 Acknowledgements

This research was conducted within the Developmental Imaging research group, Murdoch Childrens Research Institute and the Peninsula Clinical School, Monash University. It was supported by the Murdoch Childrens Research Institute, the Royal Children’s Hospital, Monash University and the Victorian Government’s Operational Infrastructure Support Program. The project was generously supported by the Royal Children’s Hospital Foundation devoted to raising funds for research at The Royal Children’s Hospital.

## A Supplementary Material

### A.1 Visualization of additional FreeSurfer estimates

#### A.1.1 Cross sectional processing, Set A

Trajectories of supratentorial volume for Set A, estimated using the cross sectional FreeSurfer pipeline are shown in Supplementary Figure S1.

#### A.1.2 Cross sectional processing, Set B

Supplementary Figures S2, S3 and S4 illustrate supratentorial volume and cortical thickness estimates from the cross sectional FreeSurfer pipeline in the presence simulated cortical and WM atrophy respectively.

**Figure S1:**
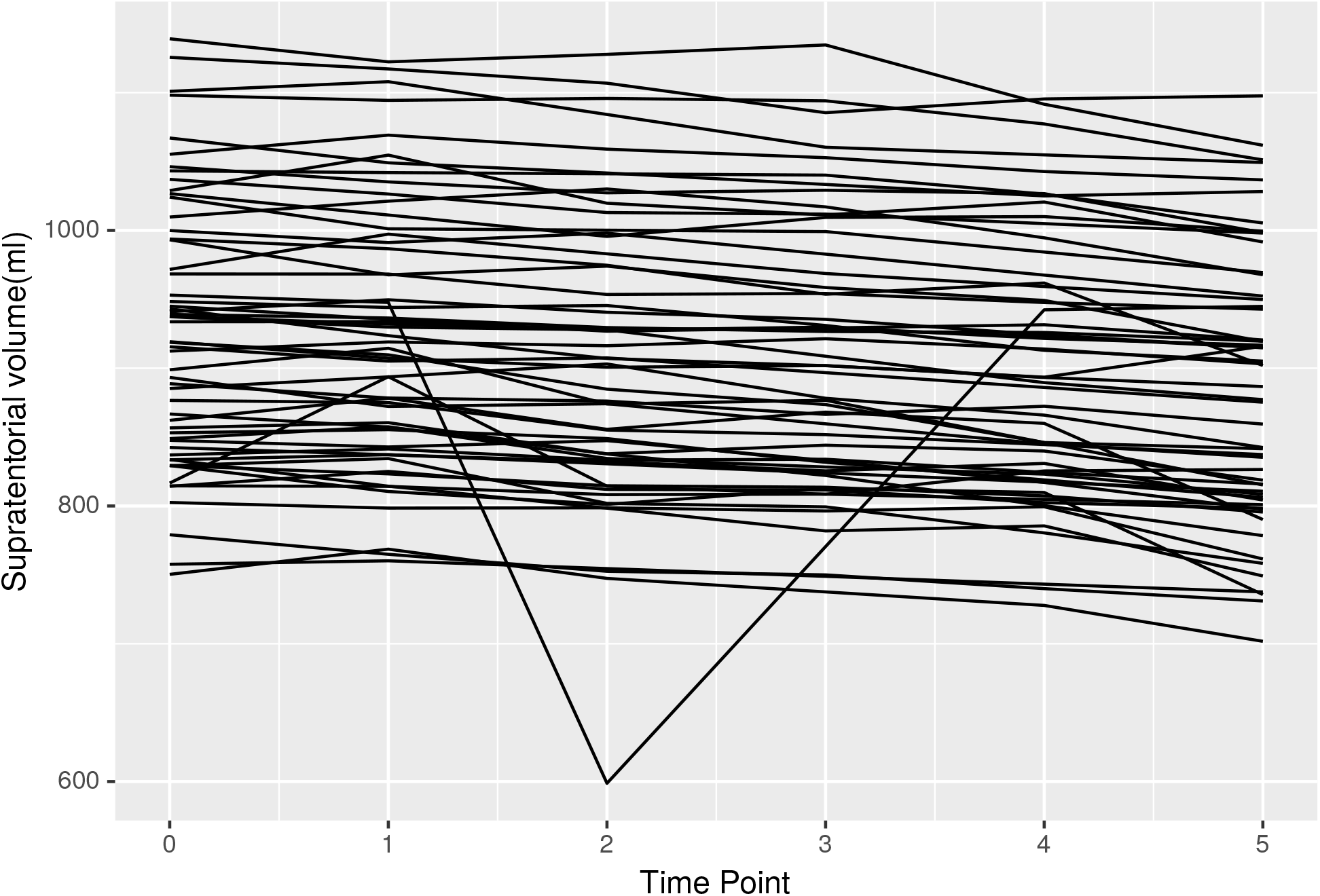
Supratentorial volume estimated with cross sectional FS for ADNI participants with scans at 60 months

**Figure S2:**
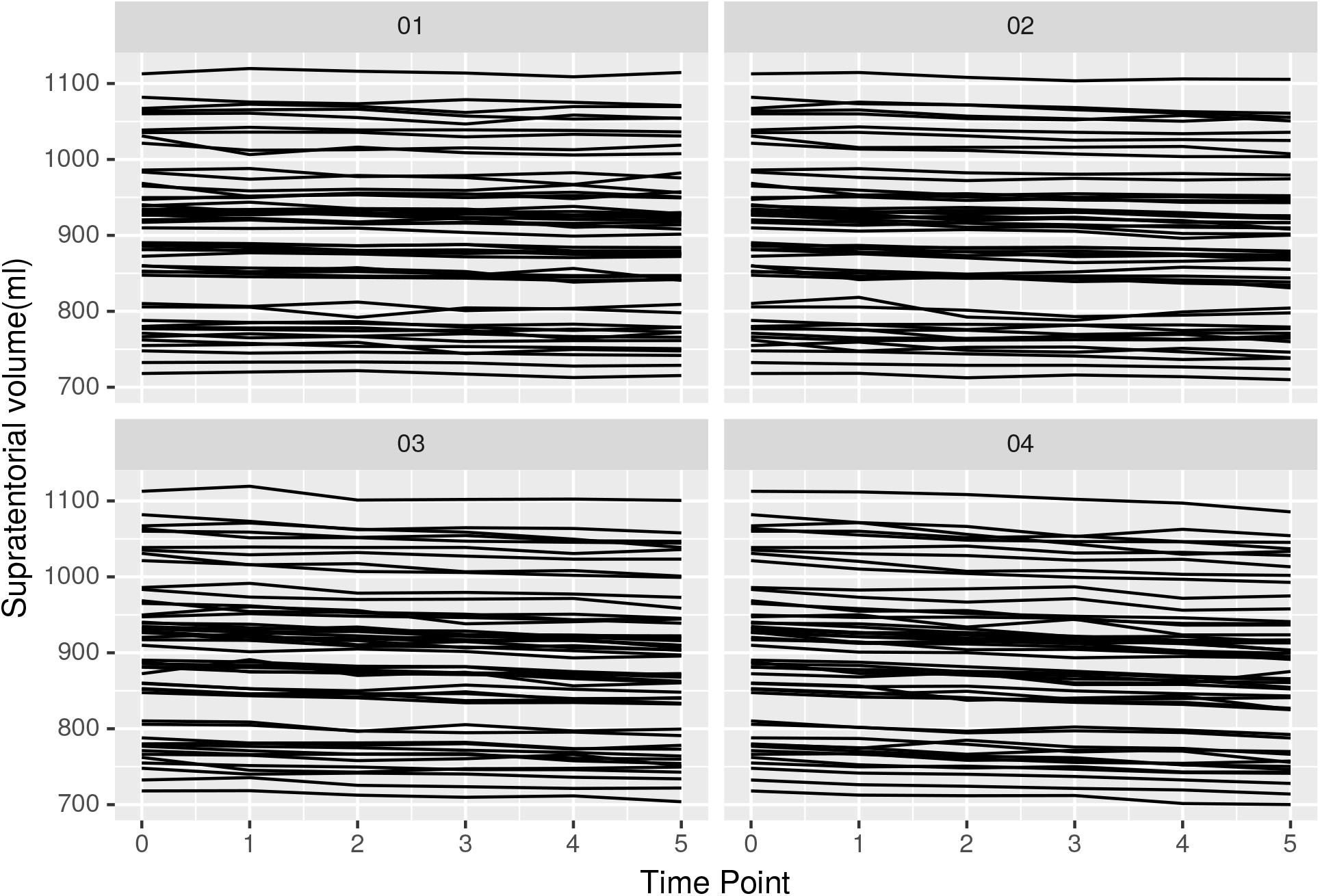
Supratentorial volume estimated with cross sectional FS in presence of simulated cortical atrophy. Each panel corresponds to a different level of simulated atrophy.

**Figure S3:**
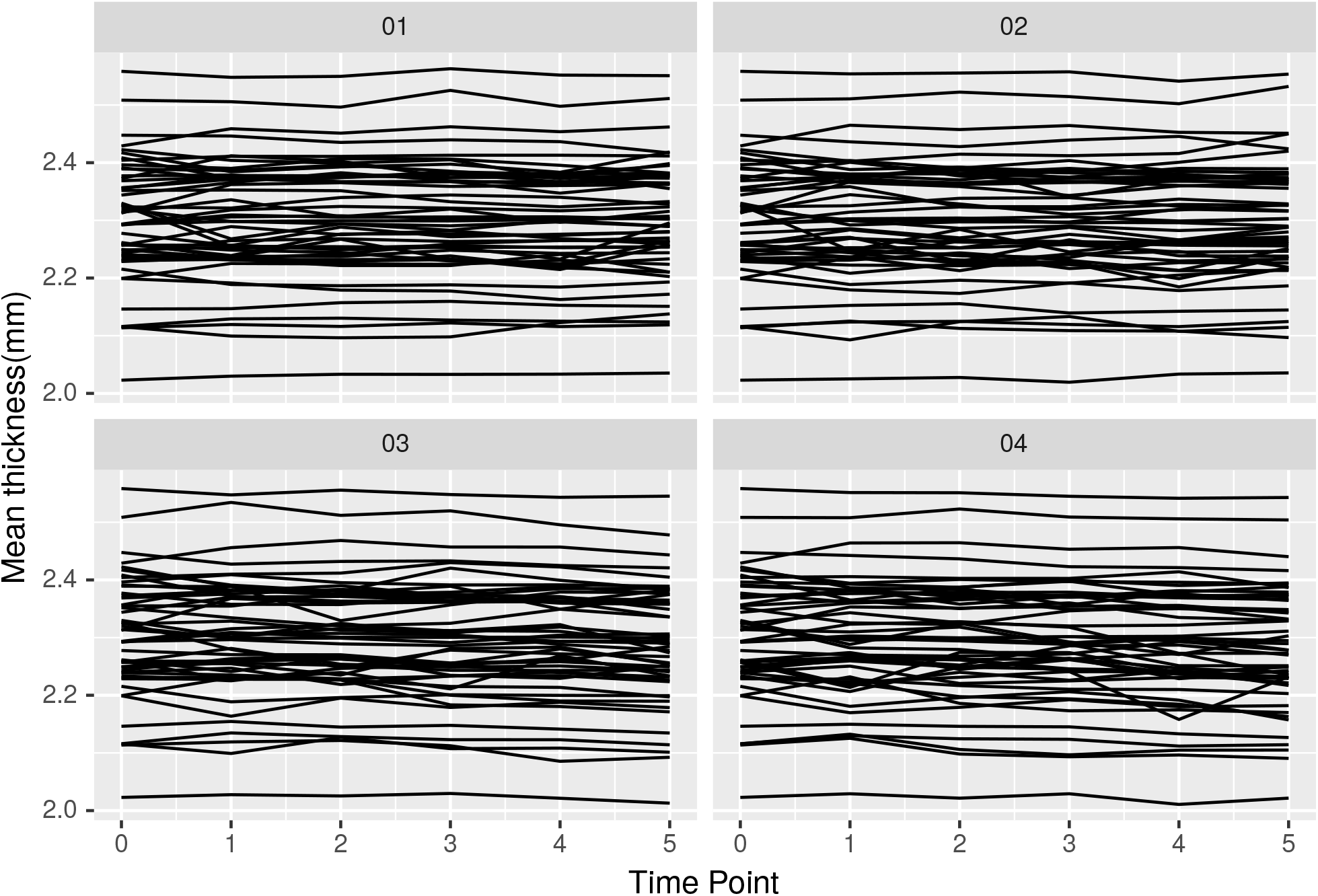
Mean cortical thickness estimated with cross sectional FreeSurfer in presence of simulated WM atrophy. Each panel corresponds to a different level of simulated atrophy.

**Figure S4:**
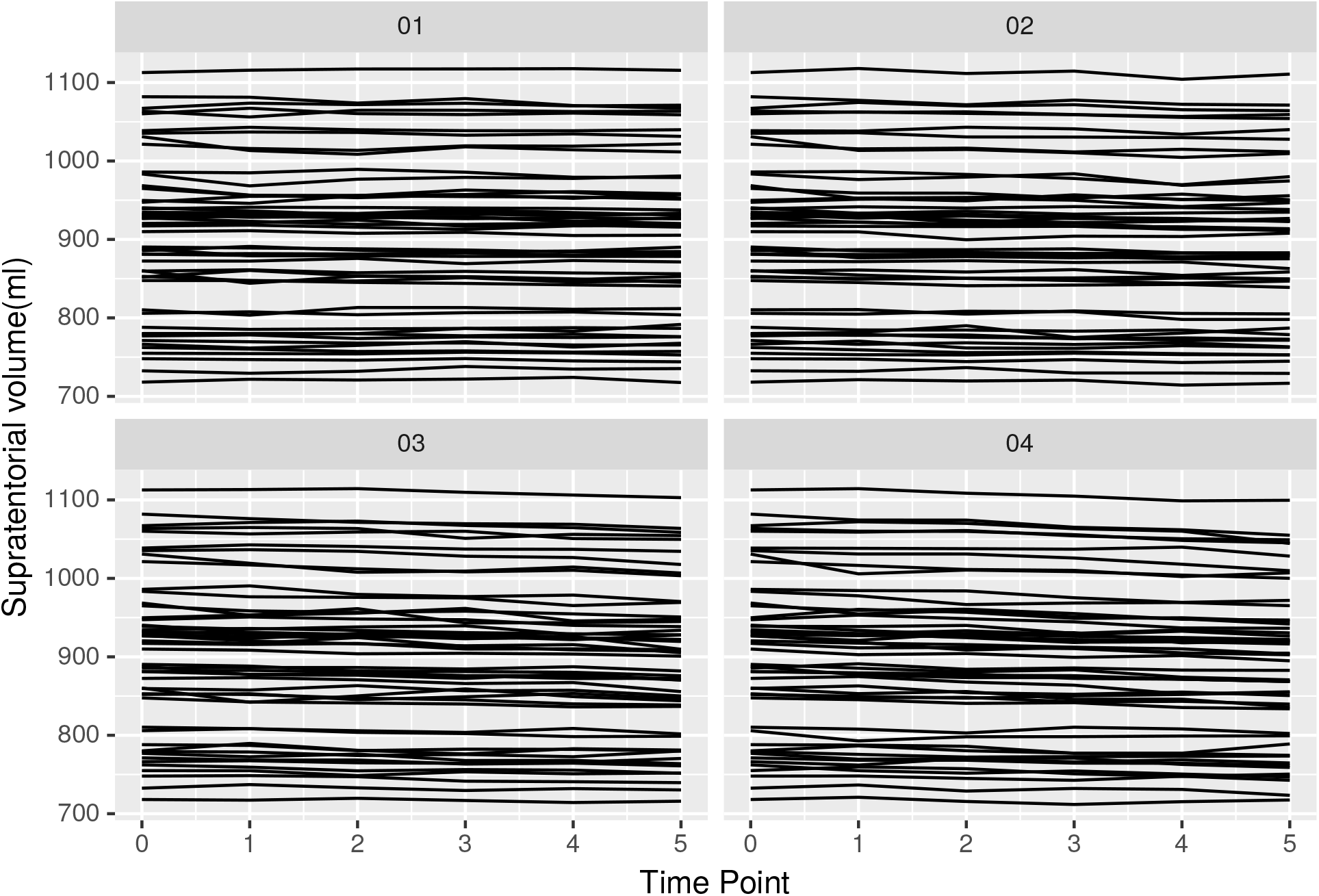
Supratentorial volume estimated with cross sectional FS in presence of simulated WM atrophy. Each panel corresponds to a different level of simulated atrophy.

#### A.1.3 Longitudinal Processing Results, Set B

Supplementary Figures S5, S7 and S6 illustrate supratentorial volume and cortical thickness estimates from the longitudinal FreeSurfer pipeline in the presence simulated cortical and WM atrophy respectively.

**Figure S5:**
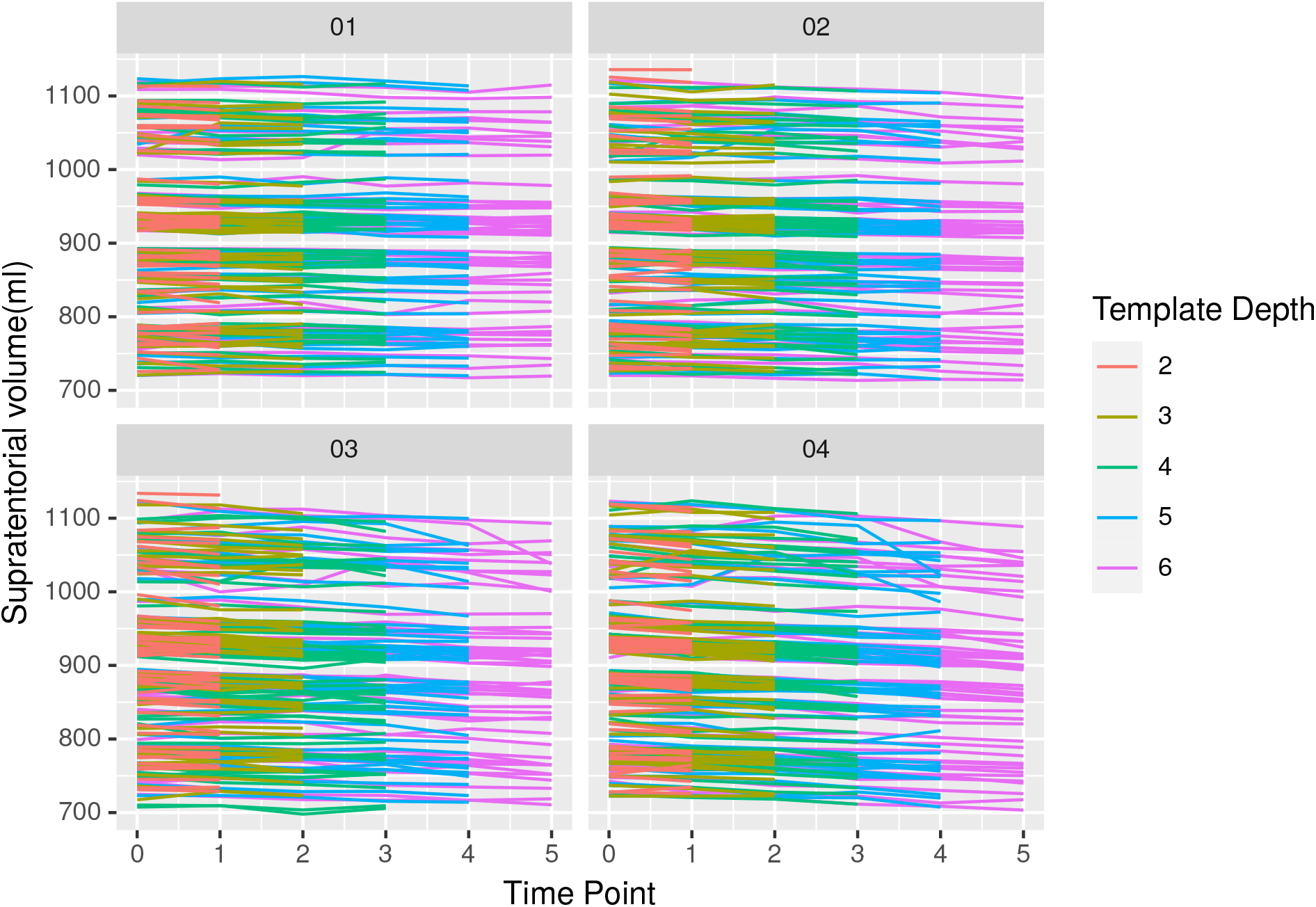
Supratentorial volume estimated with longitudinal FS in presence of simulated cortical atrophy. Each panel corresponds to a different level of simulated atrophy.

### A.2 Statistical model formulae

#### A.2.1 Group comparisons

The following lme4 formulae were used to evaluate group differences in brain measure (mean cortical thickness and supra tentorial volume). TimePoint was a continuous variable and TemplateDepthGroup was a categorical variable with two levels (for example levels corresponding to template depths of 3 and 6).

The first model was used to investigate group differences in mean, while second model, with the interaction term, was used to investigate differences in trajectory.

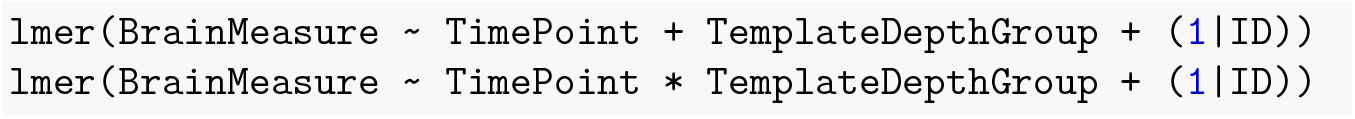

**Figure S6:**
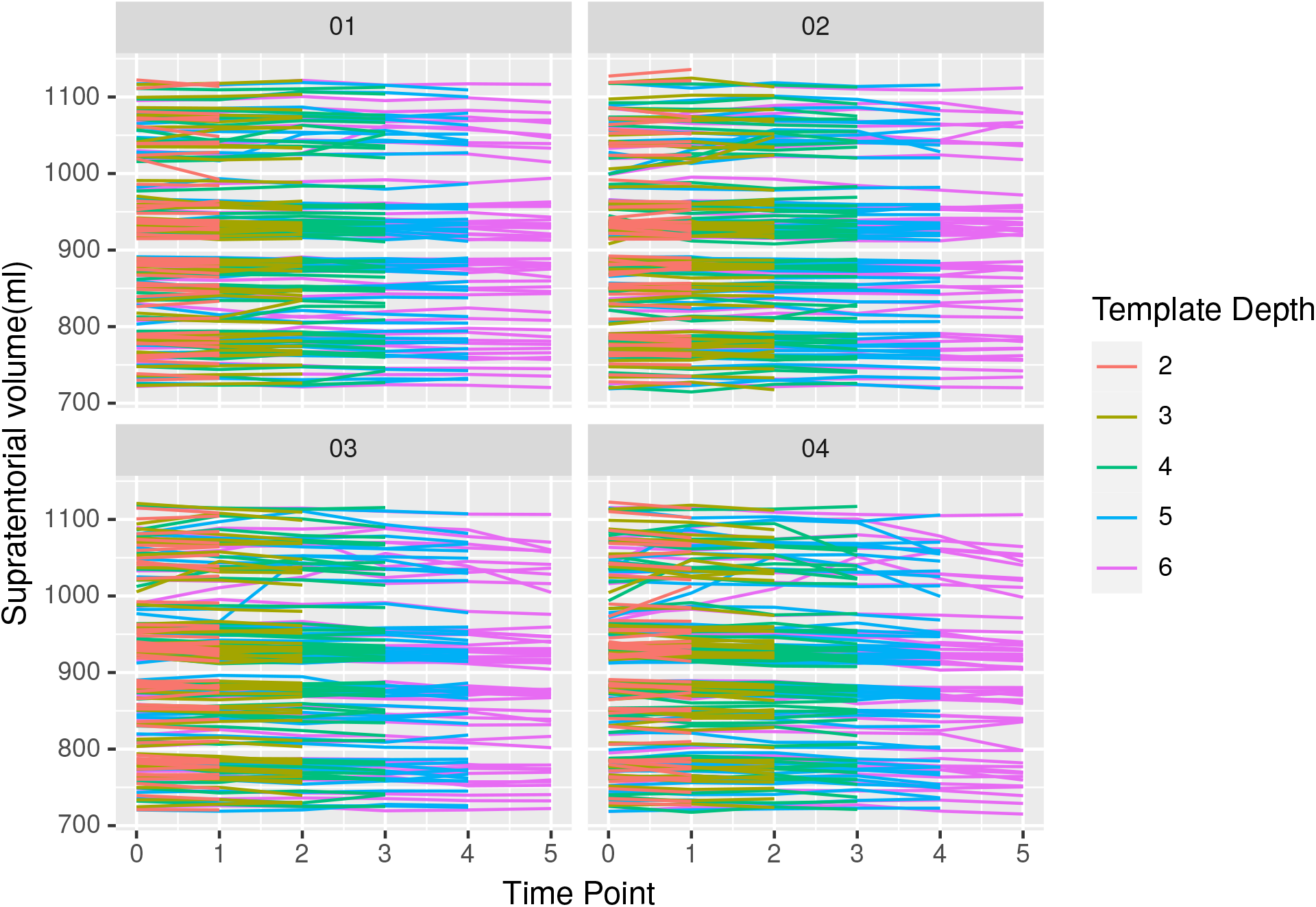
Supratentorial volume estimated with longitudinal FS in presence of simulated WM atrophy. Each panel corresponds to a different level of simulated atrophy.

**Figure S7:**
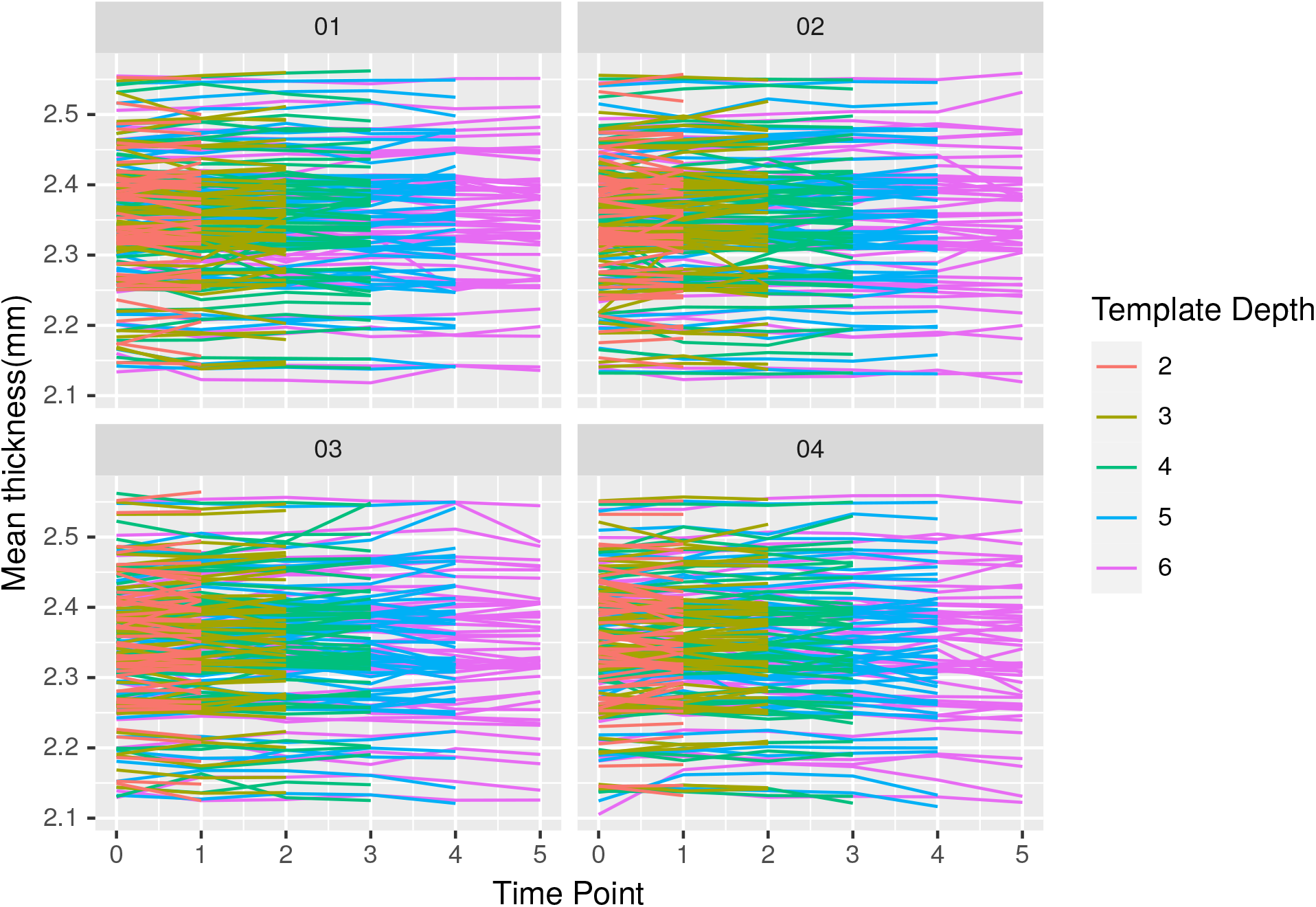
Cortical thickness estimated with longitudinal FreeSurfer pipeline and varying numbers in template in presnece of simulated WM atrophy. Each panel corresponds to a different level of simulated atrophy.

#### A.2.2 Template depth as a predictor

The formula used to investigate the relationship between template depth and mean thickness in Set A (the real ADNI data) was:

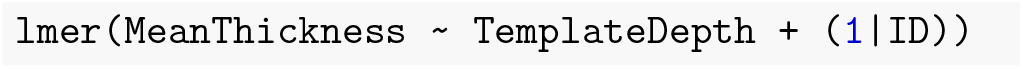

The dataset for which this model was fitted contained multiple estimates of baseline thickness, each corresponding to a different template depth, for each participant. TemplateDepth was a continuous variable

The relationship between template depth and brain measure in Set B, the simulated atrophy data, was investigated with the following models. The models were fitted separately to each simulated atrophy level and both TimePoint and TemplateDepth were continuous variables. The model with an interaction term was used to investigate the relationship between template depth and trajectory of brain measure.

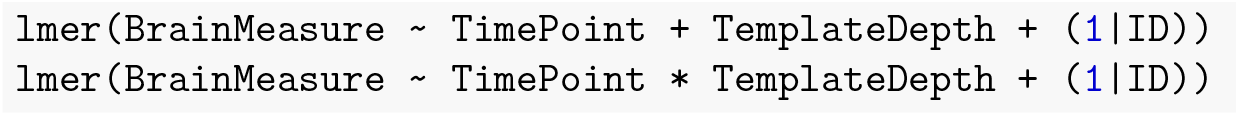

### A.3 Group comparisons for WM atrophy

#### A.3.1 Cortical thickness

**Figure S8:**
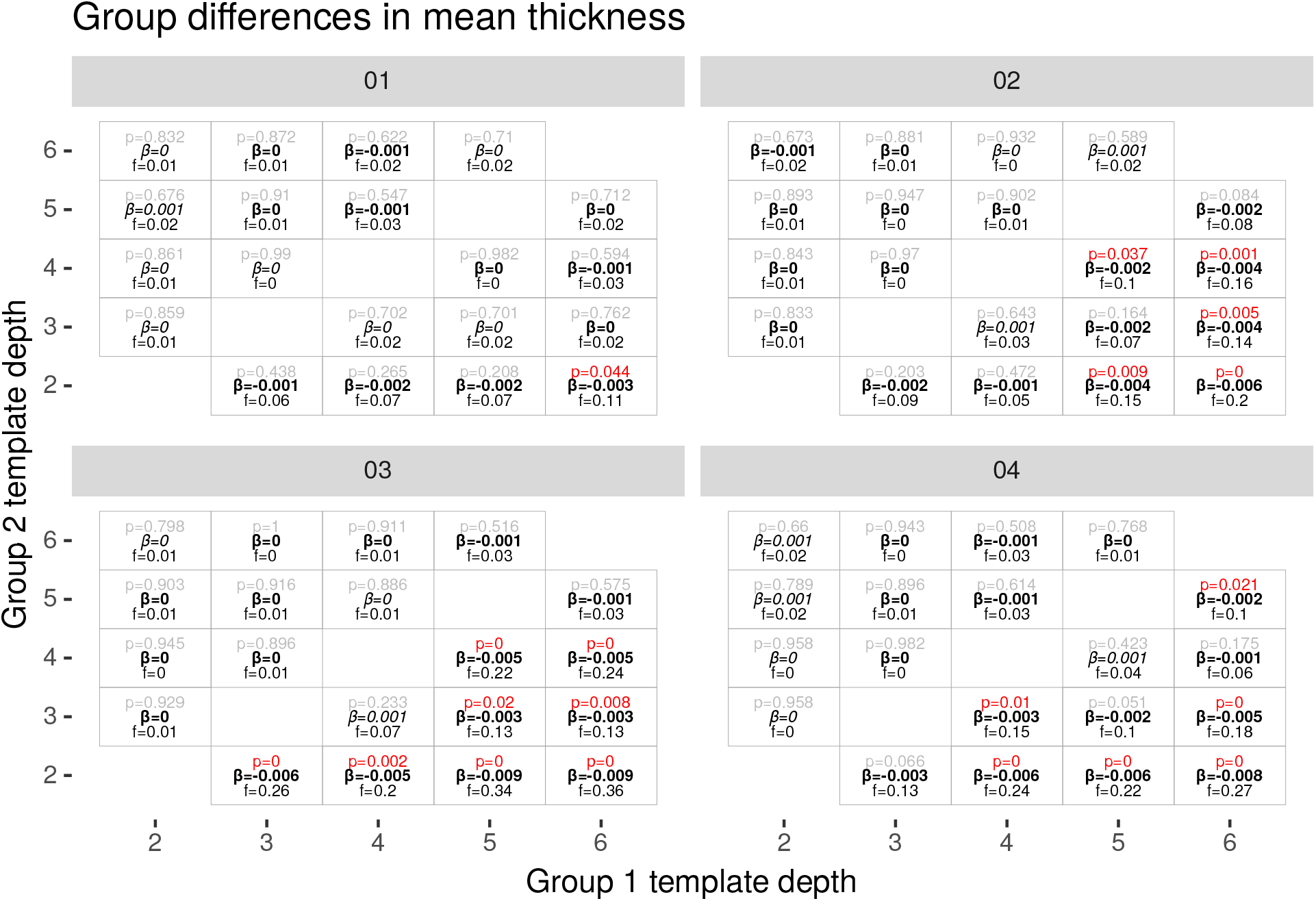
Group differences in cortical thickness for simulated WM atrophy.

Differences in mean cortical thickness between template groupings were detected in 25 of the 50 comparisons of simulated WM atrophy and the longitudinal processing pipeline (Figure S8). Minimum p value = 1e-11.

**Figure S9:**
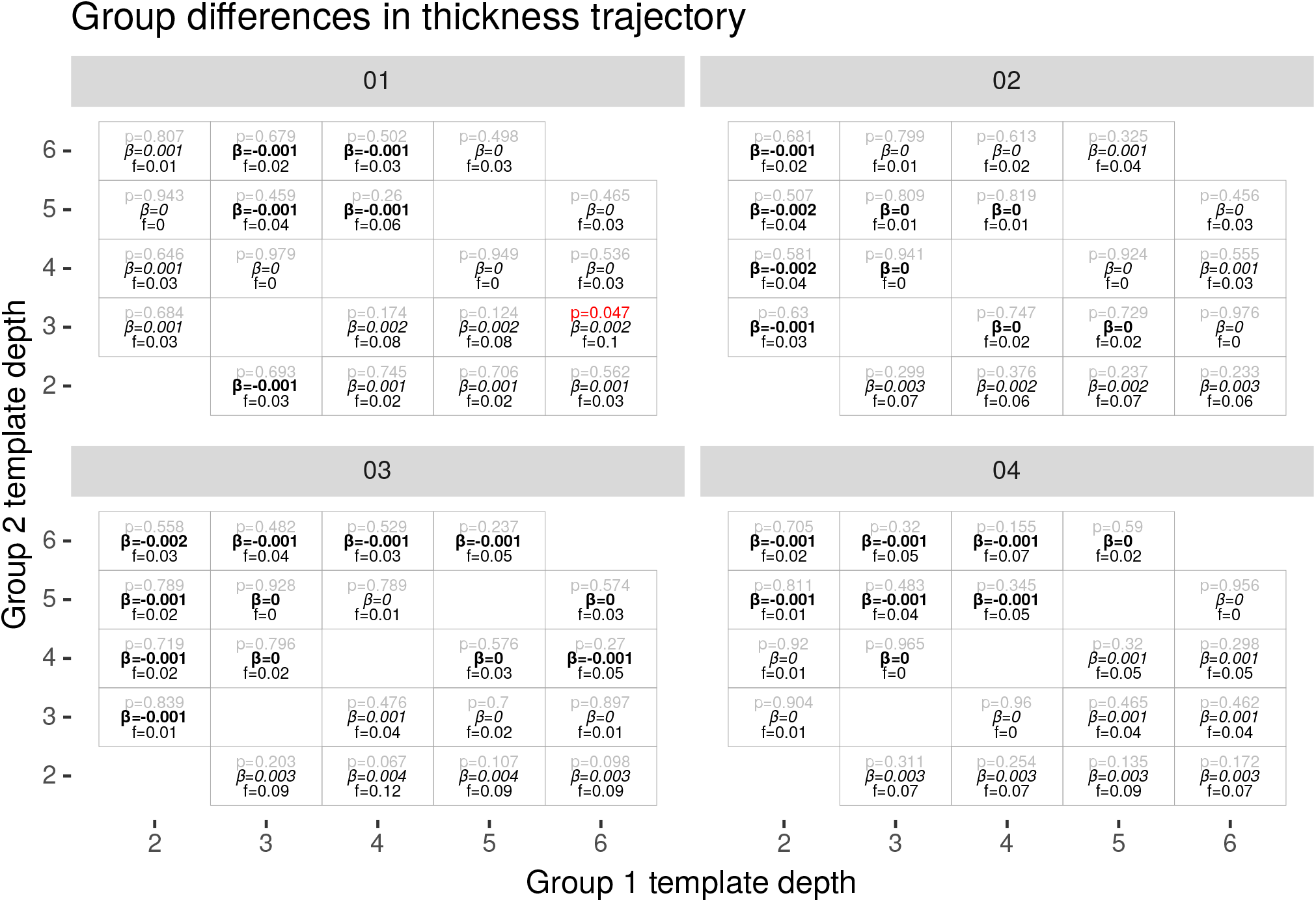
Group differences in trajectory of cortical thickness for simulated WM atrophy.

The minimum p-value (1e-11) occurred when comparing template depths 2 and 6 at atrophy level 03. Group difference in mean =-0.00882mm(stderr 0.00131) in 2.35 (Cohen’s f = 0.36)

The sign of the group difference was negative in 24 of 25 cases, meaning a lower thickness estimate for deeper templates.

Only 1 difference in trajectory detected for simulated WM atrophy (p = 0.047) occurred when comparing templates depth 3 and 6 at atrophy level 01 (Figure S9). Group difference in trajectory = 0.00229 and the slope of base group = -0.00177 (Cohen’s f = 0.10).

#### A.3.2 Supratentorial volume

Difference in mean supratentorial volume between template depth groups were detected in 20 of the 50 comparisons of simulated WM atrophy and the longitudinal processing pipeline (Figure S10).

The minimum p value (p=2.97e-5) occurred when comparing template depths 2 and 6 at atrophy level 02. The group difference in mean was -3251 (stderr 768) in 906529 (Cohen’s f = 0.23).

The sign was negative in 18 of the 20 cases, indicating lower volume for the deeper template group.

No differences in trajectory between template groups were detected for simulated WM atrophy (Supplementary Figure S11).

**Figure S10:**
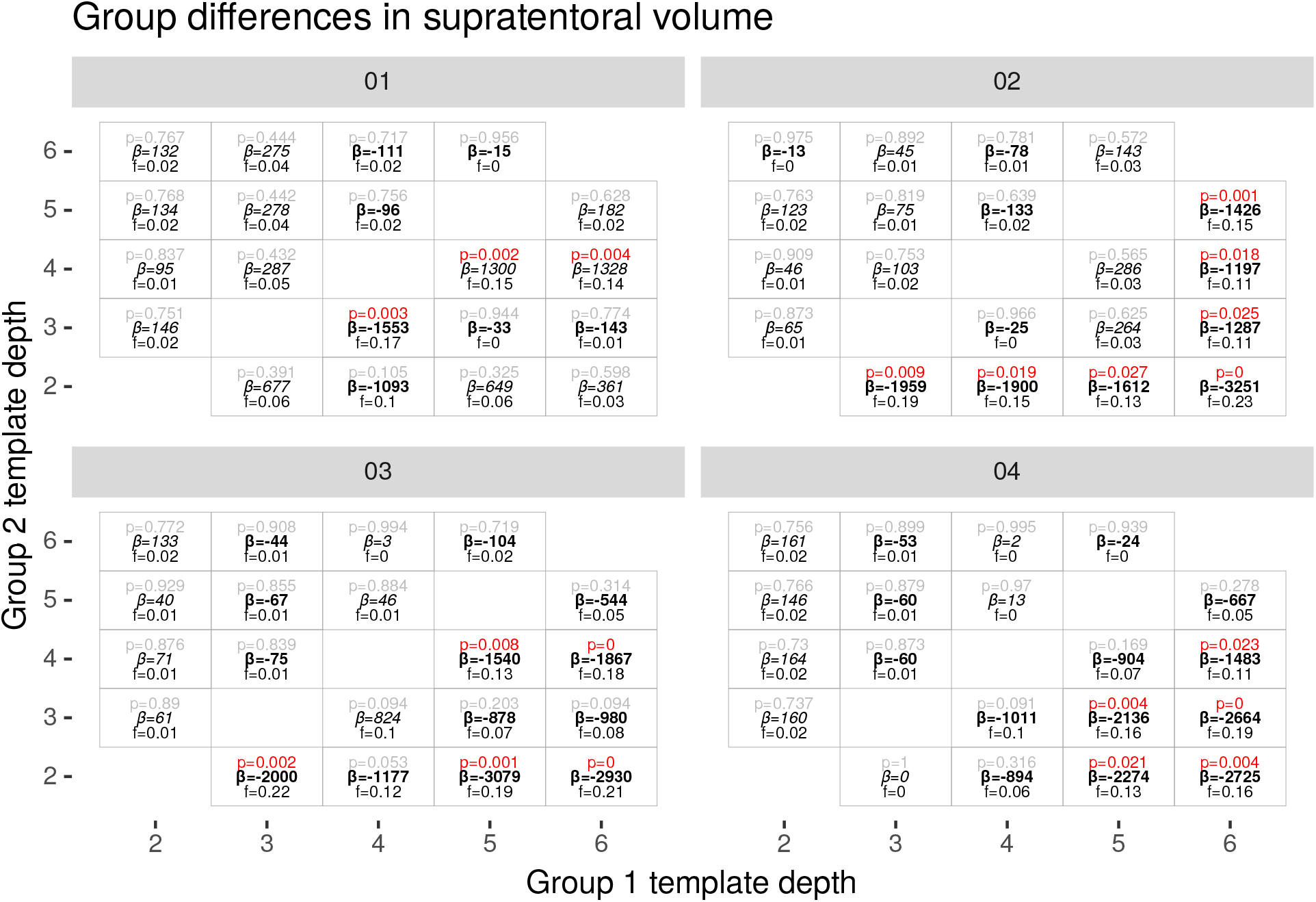
Group differences in supratentorial volume for simulated WM atrophy.

**Figure S11:**
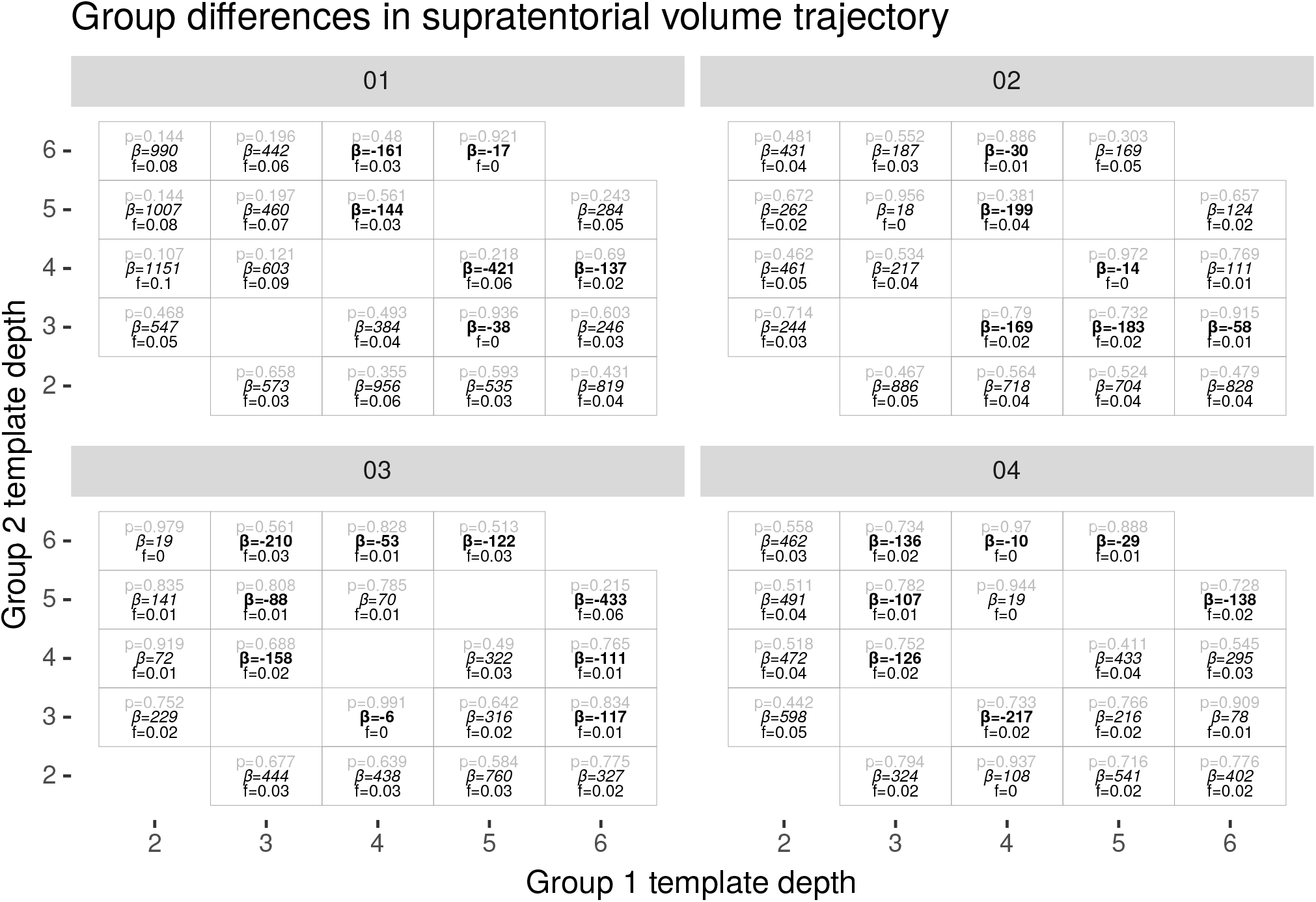
Group differences in trajectory of supratentorial volume for simulated WM atrophy.

